# Pth4 neurons define a novel hypothalamic circuit that promotes sleep via brainstem monoaminergic neurons

**DOI:** 10.1101/2025.09.09.675233

**Authors:** Ulrich Herget, Steven Tran, Chanpreet Singh, Grigorios Oikonomou, Soojin Ryu, Josep Rotllant, David A. Prober

## Abstract

Classical studies identified a critical role for the hypothalamus in regulating sleep and wake states, but few such hypothalamic neuronal populations have been identified. Here we describe a sleep-promoting population of hypothalamic neurons that expresses the neuropeptides QRFP and parathyroid hormone 4 (Pth4) in zebrafish. Optogenetic stimulation of these neurons results in a large increase in sleep that requires *pth4* but not *qrfp*. Noradrenergic locus coeruleus (LC) neurons and serotonergic raphe neurons (RN) in the hindbrain express distinct *pth receptors*, and genetic epistasis and cell ablation experiments revealed that Pth4 neuron-induced sleep is suppressed in mutants that lack noradrenaline in the LC or lack the serotonergic RN. Pth4 neuron-induced sleep is also suppressed in *serine/threonine kinase 32a* (*stk32a*) mutants, possibly via *stk32a*-expressing neurons in the prethalamus that express *pth receptors*. These results identify QRFP/Pth4 neurons as a novel hypothalamic sleep-promoting population and support a model in which distinct sleep- and wake-promoting hypothalamic populations act via monoaminergic neurons in the hindbrain to control vigilance state.

## Introduction

Sleep occupies a third of the human lifespan, is universal among animals, and is essential for life, but the genetic and neuronal mechanisms that regulate sleep remain poorly understood ^1–4^. Classical studies identified the hypothalamus as a brain structure that plays a critical role in regulating sleep ^5^, and more recent studies have identified distinct sleep- and wake-promoting populations in the hypothalamus ^6^. For example, the discovery of wake-promoting neurons that express the neuropeptide hypocretin/orexin (Hcrt) in the lateral hypothalamus in humans ^7,8^, rodents ^9,10^, and zebrafish ^11–13^ identified an important and evolutionarily conserved source of arousal. A small number of sleep-promoting hypothalamic neurons and neuropeptides have also been identified. For example, melanin-concentrating hormone (MCH) and MCH-expressing neurons promote sleep in rodents ^14–18^ and zebrafish ^19^, and hypothalamic neurons that express neuropeptide VF (NPVF) promote sleep in zebrafish ^20,21^. However, the mechanisms through which these neurons interact with each other, and with downstream neural circuits in the brain, to regulate sleep/wake states are poorly understood. It is also likely that additional sleep-promoting populations remain to be discovered.

The zebrafish has emerged as a useful vertebrate model to discover genes and neurons that regulate sleep, and to determine the mechanisms through which they act ^22–24^. Larval zebrafish have several useful features for this purpose, including a small and transparent brain that is anatomically and molecularly similar to the mammalian brain ^25,26^, and amenability to large-scale screens ^27,28^ and high-throughput sleep assays ^13^. Drug screens ^27,29^ and targeted perturbations of specific genes and neurons ^13,19,30–32^ have shown that most mechanisms that affect mammalian sleep have similar effects on larval zebrafish behavior, suggesting that most mechanisms that regulate mammalian sleep are conserved in larval zebrafish. We previously used a candidate-based approach to identify a population of hypothalamic neurons that express the neuropeptide QRFP ^33,34^ (also known as P518 and 26RFa) ^35–40^, and showed that overexpression of QRFP had a mildly suppressive effect on locomotor activity without affecting sleep, while loss of *qrfp* or its receptors *gpr103a* and *gpr103b* resulted in a small decrease in sleep during the day ^34^. These observations suggested that QRFP signaling plays a minor role, if any, in regulating sleep.

Here we explore the role of *qrfp*-expressing neurons in zebrafish sleep, and show that stimulation of these neurons potently induces sleep. Surprisingly, we found that *qrfp* is dispensable for this phenotype. Instead, we discovered that the sedating effect of these neurons requires *parathyroid hormone 4* (*pth4*), a neuropeptide that is exclusively co-expressed with *qrfp* in the zebrafish hypothalamus, and was previously shown to regulate bone remodeling ^41,42^, but had not been implicated in regulating sleep. We show that QRFP/Pth4 neurons promote sleep via interactions with the wake-promoting noradrenergic (NA) locus coeruleus (LC) ^31^, the sleep-promoting serotonergic (5HT) raphe nuclei (RN) ^32^, and the recently identified sleep regulator *serine/threonine kinase 32a* (*stk32a*) (Tran et al., submitted), likely via distinct Pth receptors. This study thus identifies a novel population of sleep-promoting neurons in the hypothalamus, and reveals that these neurons promote sleep via monoaminergic populations in the hindbrain that are known to regulate sleep in both zebrafish and mice. Together with other studies ^3,20,31,43,44^, our results support a model in which distinct hypothalamic peptidergic neurons promote sleep or wake states by activating or inhibiting specific monoaminergic populations in the hindbrain. These hindbrain populations thus act as hubs that integrate distinct arousing and sedating signals to determine vigilance state.

## Results

### Hypothalamic QRFP neurons promote sleep

Previous experiments showed that overexpression of QRFP results in a decrease in locomotor activity during the day, with no effect on sleep, whereas a predicted null mutation in *qrfp* results in a small increase in locomotor activity and a small decrease in sleep during the day ^34^. Thus, QRFP signaling can suppress locomotor activity and promote sleep, but the magnitudes of these phenotypes are small. QRFP is exclusively expressed in the hypothalamus of larval zebrafish ^33,34^ (21 ± 1 neurons total in both brain hemispheres, n=23 fish at 5 days post-fertilization (dpf)). These neurons are rostral to and partially comingled with, but distinct from, neurons that express the wake-promoting neuropeptide Hcrt (Figures 1A and 1A’, Video S1) (1/294 *qrfp* neurons co-express *hcrt*, n=13 fish) ^11–13^, and are also distinct from neurons that express the sleep-promoting neuropeptide NPVF (Figures 1C and 1C’, Video S2) (0/195 *qrfp* neurons co-express *npvf*, n=10 fish) ^20,21,45^. This region of the zebrafish brain corresponds to the mammalian lateral hypothalamus, where these three neuropeptides are also expressed ^9,46–48^. These three neuronal populations project to both shared and distinct targets (see below), suggesting they may affect sleep via at least some shared postsynaptic targets.

**Figure 1.**
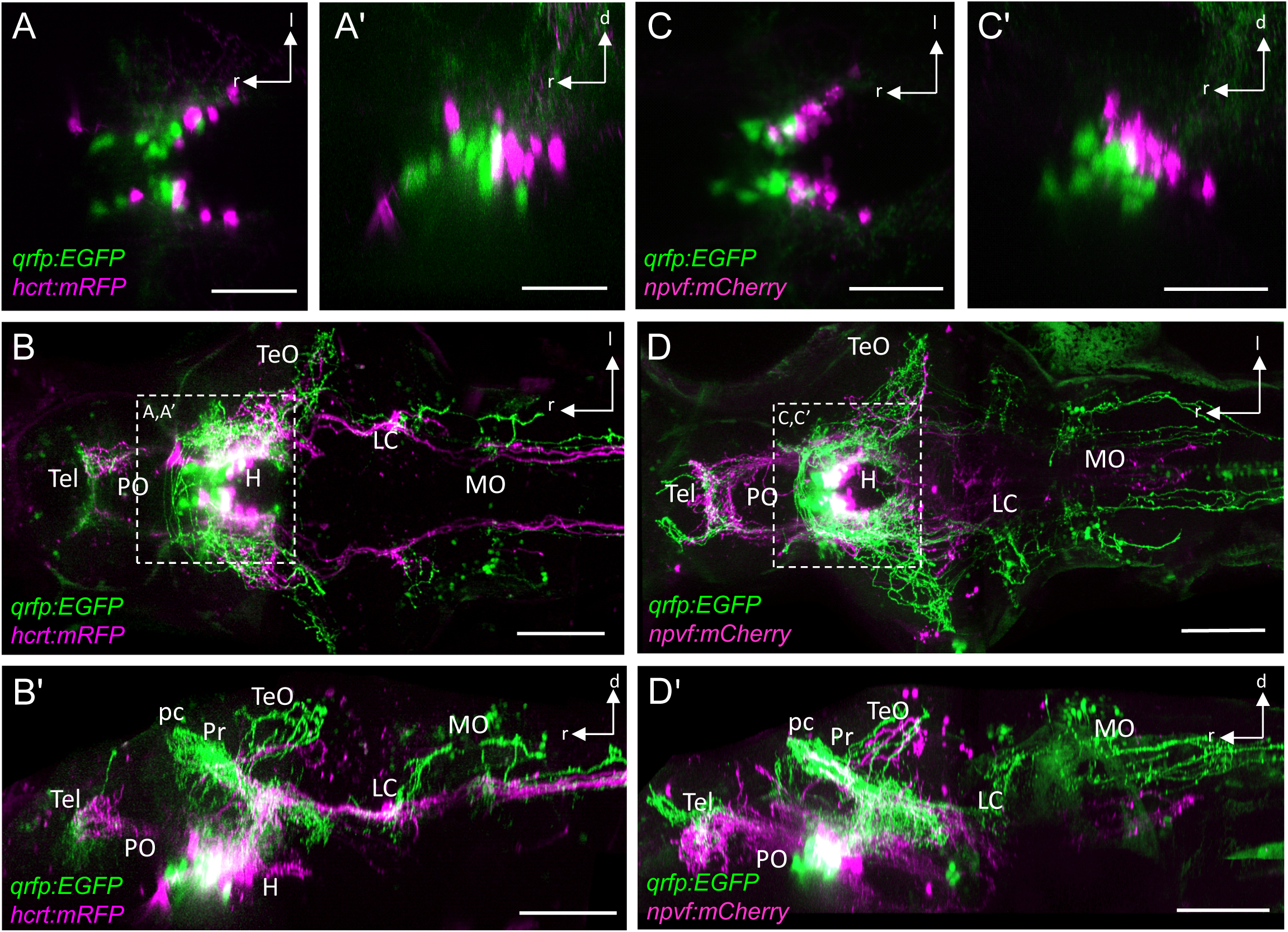
QRFP, Hcrt, and NPVF are expressed in distinct neuronal populations in the larval zebrafish hypothalamus. Dorsal (**A,B,C,D**) and lateral (**A’,B’,C’,D’**) maximum intensity projection views of live 5-dpf zebrafish that express EGFP in QRFP neurons and mRFP in Hcrt neurons (**A-B’**), or mCherry in NPVF neurons (**C-D’**). (**A,A’,C,C’**) Substacks showing hypothalamic somata. (**B,B’,D,D’**) Whole brain stacks with increased signal intensity to show neuronal projections. Boxed regions in (**B,D**) are shown with reduced signal intensity in (**A,A’,C,C’**). l, lateral; d, dorsal; r, rostral; H, hypothalamus; LC, locus coeruleus; MO, medulla oblongata; pc, posterior commissure; Pr, pretectum; PO, preoptic area; Tel, telencephalon; TeO, optic tectum. Scale bars: 50 μm (**A,A’,C,C’**), 100 μm (**B,B’,D,D’**).

Prior zebrafish studies showed that optogenetic stimulation of Hcrt neurons promotes wakefulness ^31^, while stimulation of NPVF neurons promotes sleep ^20^. In order to explore the function of QRFP neurons, we generated transgenic zebrafish in which these neurons express ReaChR (Figure S1) ^49^, and stimulated these neurons using blue light while monitoring the behavior of 96 freely-swimming fish. We found that stimulating QRFP neurons resulted in a large decrease in locomotor activity and increase in sleep during both the night and day (Figure 2), indicating that stimulation of QRFP neurons results in increased sleep.

**Figure 2.**
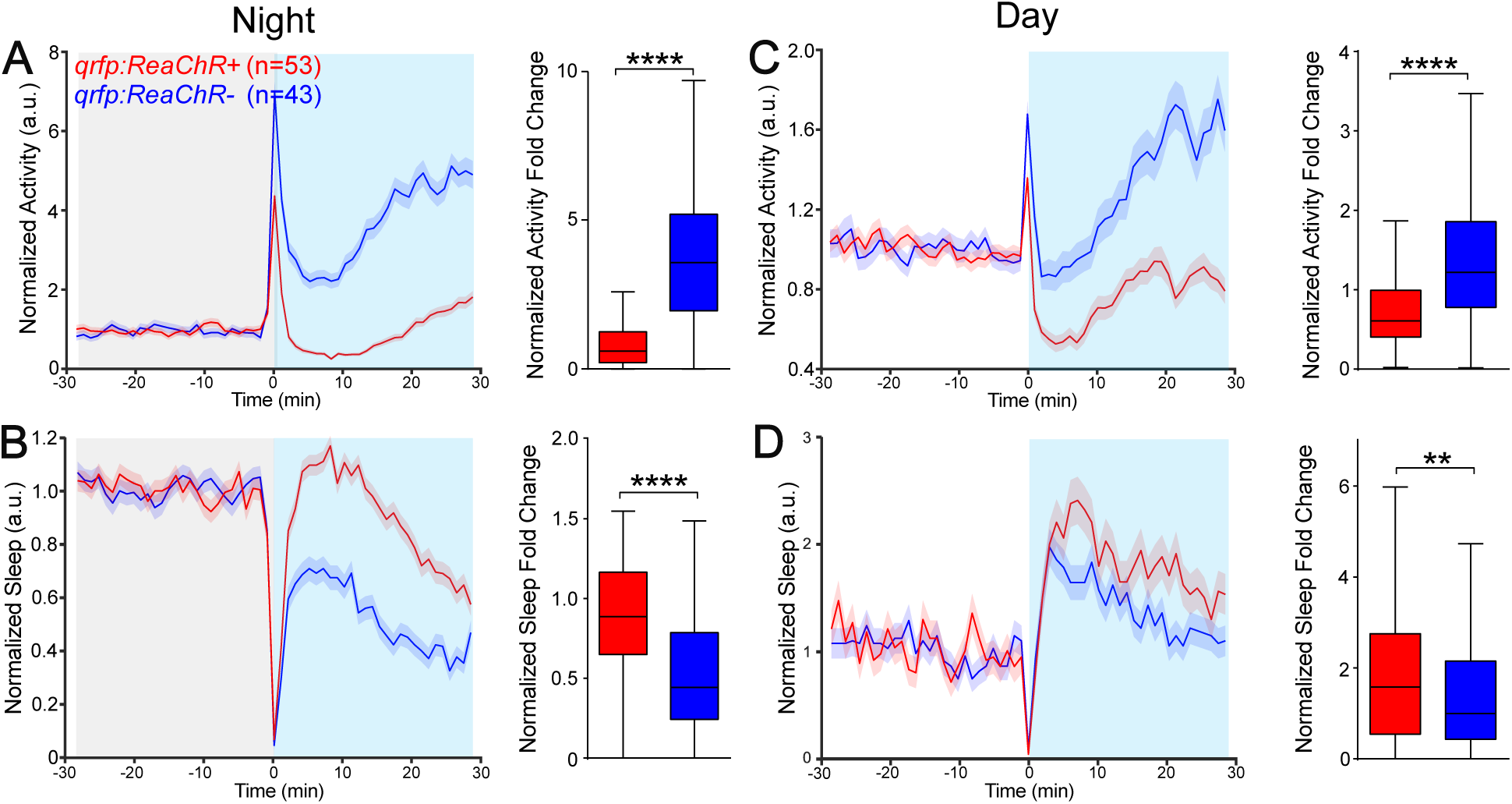
Optogenetic stimulation of QRFP neurons suppresses locomotor activity and increases sleep. Fish in which *qrfp*-expressing neurons are optogenetically stimulated (red) show a decrease in locomotor activity (**A,C**) and an increase in sleep (**B,D**) compared to non-transgenic sibling controls (blue) both at night (**A,B**) and during the day (**C,D**). Gray and blue shading indicate darkness and blue light exposure, respectively. Values during blue light exposure are normalized to values during 30 minutes prior to blue light exposure. Blue light onset induces a brief startle response which is excluded from the analysis. Mean ± SEM line plots and Tukey box plots are shown. n = number of fish. **, p<0.01, ****, p<0.0001 by Mann-Whitney U test.

### QRFP neuron-induced sleep requires *pth4* but not *qrfp*

Based on previous zebrafish QRFP neuropeptide gain- and loss-of-function observations ^34^, we hypothesized that the ability of QRFP neurons to promote sleep depends on the QRFP neuropeptide. To test this hypothesis, we optogenetically stimulated QRFP neurons in animals that were either heterozygous or homozygous for a predicted null mutation in the *qrfp* gene ^34^. Surprisingly, the sedating effect of stimulating QRFP neurons was unaffected in *qrfp* -/- fish compared to their *qrfp* +/- siblings (Figures 3A-3D), suggesting that these neurons promote sleep via a different mechanism.

**Figure 3.**
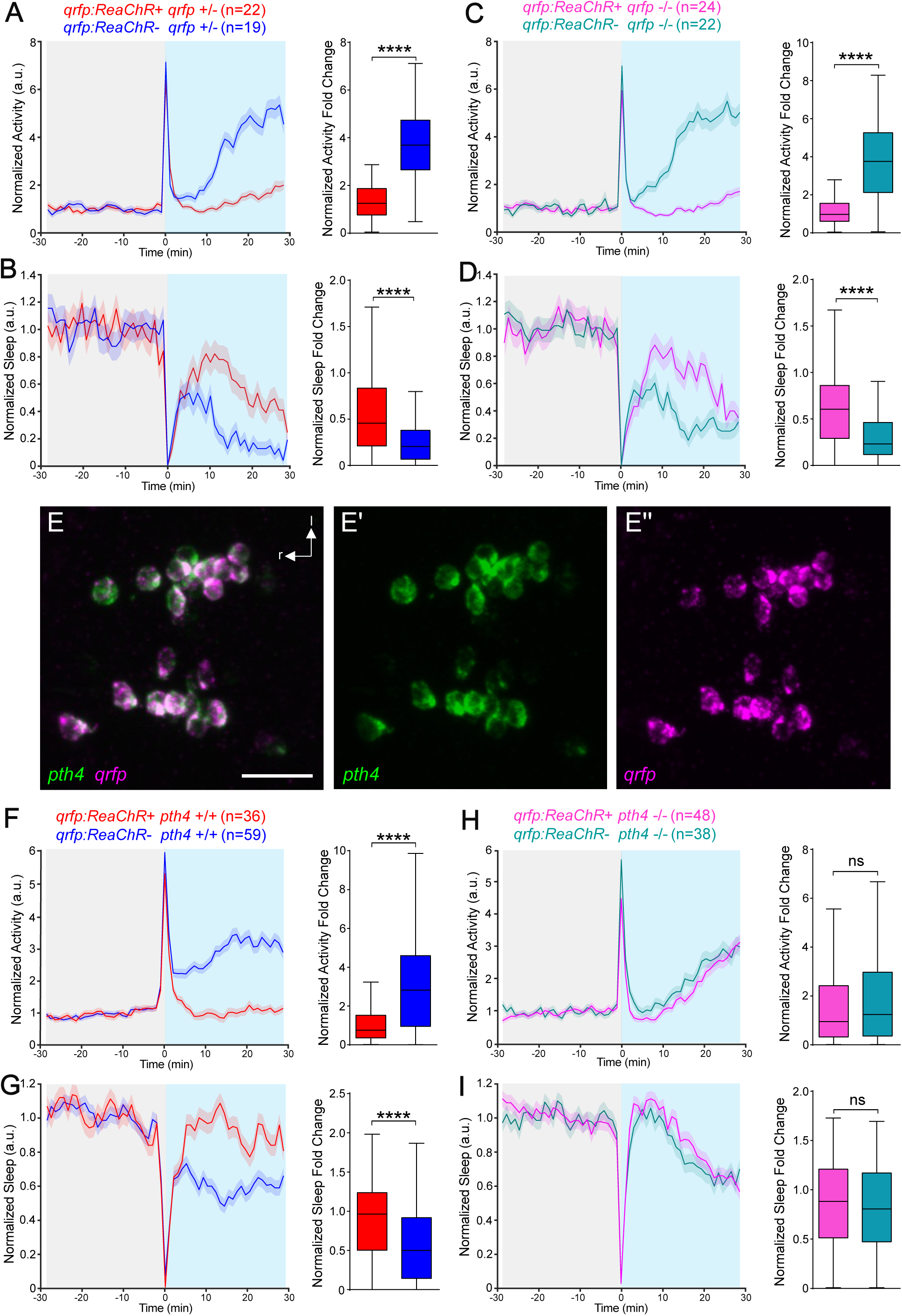
Sleep induction by optogenetic stimulation of QRFP/Pth4 neurons requires *pth4* but not *qrfp*. (**A-D**) Optogenetic stimulation of QRFP/Pth4 neurons suppresses locomotor activity and increases sleep in *qrfp* +/- (**A,B**) and *qrfp* -/- fish (**C,D**) compared to non-transgenic siblings. (**E**) FISH shows that *pth4* and *qrfp* are co-expressed in the same hypothalamic neurons (586/589 cells, n=23 fish). Scale bar: 25 μm. (**F-I**) Optogenetic stimulation of QRFP/Pth4 neurons suppresses locomotor activity and increases sleep in *pth4* +/+ fish (**F,G**) but not in *pth4* -/- fish (**H,I**) compared to non-transgenic siblings. Mean ± SEM line plots and Tukey box plots are shown. n = number of fish. ****, p<0.0001, ns, p>0.05 by Mann-Whitney U test.

A candidate protein for this function is the neuropeptide parathyroid hormone 4 (Pth4), as it has an expression pattern similar to that of QRFP ^41^. Using fluorescent *in situ* hybridization (FISH), we found that *pth4* and *qrfp* are co-expressed in the same neurons in the zebrafish hypothalamus (Figure 3E, 586/589 (99.5%) of Pth4 neurons co-express *qrfp*, n=23 fish), and are not expressed in any other cells in the animal. To test whether the sedating function of these neurons requires *pth4*, we generated fish with a predicted null mutation in the *pth4* gene and compared the effect of optogenetic stimulation of QRPF/Pth4 neurons in *pth4* -/- and *pth4* +/+ animals. We found that the sedating effect of stimulating QRFP/Pth4 neurons was abolished in *pth4* -/- fish (Figures 3F-3I), indicating that *pth4* is required for the sedating function of these neurons. These experiments thus identify Pth4 as a novel sleep-promoting neuropeptide, and we now refer to these neurons as *pth4*-expressing neurons.

We next tested whether Pth4 neurons are required for normal sleep levels by generating transgenic fish that express nitroreductase 2.0 (NTR 2.0) ^50^ in these neurons. NTR converts the inert pro-drug metronidazole (MTZ) into a cell-autonomous toxin, such that treating zebrafish with MTZ specifically ablates NTR-expressing cells. We found that treating these animals with 4 mM MTZ resulted in the loss of Pth4 neurons, but had little or no effect on locomotor activity or sleep compared to identically treated non-transgenic siblings (Figure S2). Similarly, fish containing a predicted null mutation in *pth4*, or both *qrfp* and *pth4,* lacked sleep phenotypes compared to sibling controls (data not shown). Based on these observations, we conclude that redundant sleep-promoting pathways likely compensate for the loss of *pth4* and *pth4*-expressing neurons to maintain normal sleep levels.

### QRFP, Hcrt and NPVF neurons project to both common and distinct brain regions

In order to explore mechanisms through which Pth4 neurons promote sleep, we first compared their projection pattern to that of other hypothalamic neurons that regulate sleep. To do so, we labeled QRFP, Hcrt, and NPVF neurons with EGFP, mRFP, and mCherry, respectively, and imaged these neurons at 5 dpf.

We found that QRFP/Pth4 neurons heavily innervate the hypothalamus, posterior commissure, pretectum, optic tectum (TeO), LC, and lateral medulla oblongata (MO), sparsely innervate the rostral telencephalon, and project down the spinal cord (Figures 1B, 1B’, 1D and 1D’). Hcrt neurons also heavily innervate the hypothalamus, and project to the caudal telencephalon, LC, lateral MO, and down the spinal cord (Figures 1B and 1B’). Thus, while the Pth4 and Hcrt neuron projection patterns are largely distinct, both populations innervate the LC, a population of noradrenergic neurons that is required for the wake-inducing function of Hcrt neurons in both zebrafish ^31^ and mice ^51^. These observations suggest that Hcrt and Pth4 neurons might induce wake and sleep, respectively, via opposite effects on the LC.

We observed that NPVF neurons heavily innervate the hypothalamus, ventral and caudal telencephalon, TeO, inferior raphe ^21^, and the medial MO, and project down the spinal cord (Figures 1D and 1D’). Thus, other than the hypothalamus and optic tectum, there is little apparent overlap in the projection patterns of Pth4 and NPVF neurons in the brain, suggesting these neuronal populations may promote sleep via distinct mechanisms.

### Pth4 receptors are expressed in neuronal populations that regulate sleep

Pth4 is thought to signal via a family of Pth receptors ^52^, of which the zebrafish genome contains four paralogs: *pth1r* ^53^, *pth2ra* ^54^, *pth2rb*, and *pth3r* ^53^. To identify neurons through which Pth4 might promote sleep, we performed hybridization chain reaction (HCR) with a tyramide amplification step to increase signal intensity (Singh et al, manuscript in preparation) to determine the expression pattern of each gene (Figure 4). We found that *pth1r* (Figures 4A-4A”) is broadly expressed in the retina and many brain regions, including the telencephalon, hypothalamus, TeO, MO, RN (see also Figures 6B and 6C), near the LC (see also Figure S3A), and in the spinal cord and developing vertebrae. *pth2ra* (Figures 4B-4B”) is expressed in much of the telencephalon, hypothalamus, and TeO, as well as near the LC (see also Figure S3B), in the prethalamus (see also Figures 7A-7A’’’) and in the MO and spinal cord. *pth2rb* (Figures 4C-C”) is expressed in the telencephalon, hypothalamus, LC (see also Figure 5C), prethalamus (see also Figures 7B-7B’’), MO, TeO, and near the superior raphe (see also Figure S4B). In contrast to the widespread expression of these receptors, *pth3r* expression (Figures 4D-4D”) is restricted to the RN (see also Figures 6D and 6E). All four receptors might participate in Pth4 signaling, as Pth4 treatment can induce signaling in human embryonic kidney cells that heterologously express *pth1r*, *pthr2ra*, or *pth3r*, and human PTH1 can induce signaling via zebrafish *pth1r* as efficiently as zebrafish Pth4^41^.

**Figure 4.**
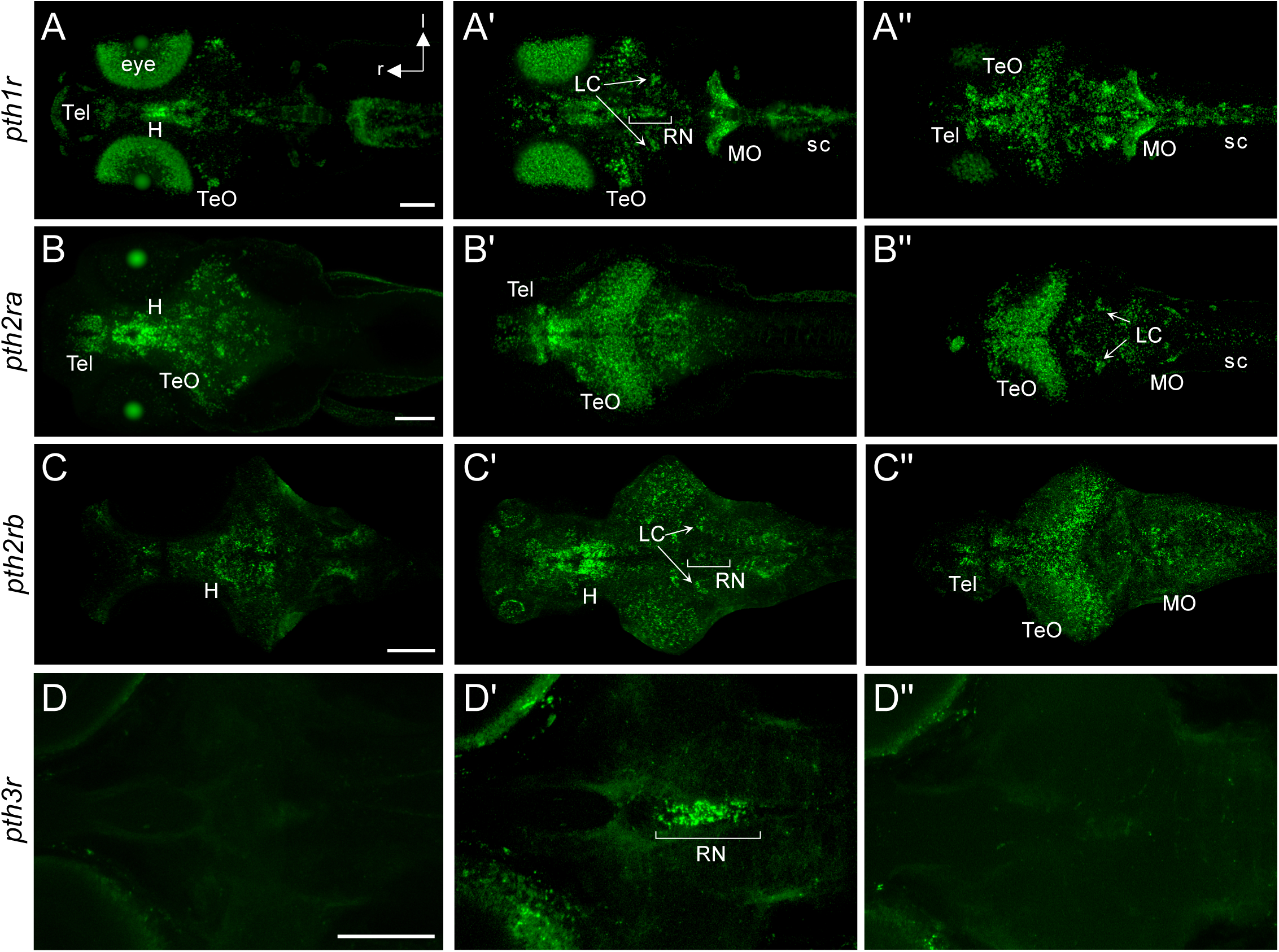
Expression of *pth receptors* in the larval zebrafish brain. HCR staining of 6 dpf larval zebrafish shows the expression patterns of *pth1r* (**A-A’’**), *pth2ra* (**B-B’’**), *pth2rb* (**C-C’’**), and *pth3r* (**D-D’’**) in ventral (**A-D**), central (**A’-D’**), and dorsal (**A’’-D’’**) substacks. l, lateral; r, rostral; H, hypothalamus; LC, locus coeruleus; MO, medulla oblongata; RN, raphe nuclei; sc, spinal cord; Tel, telencephalon; TeO, optic tectum. Scale bars: 100 μm.

**Figure 5.**
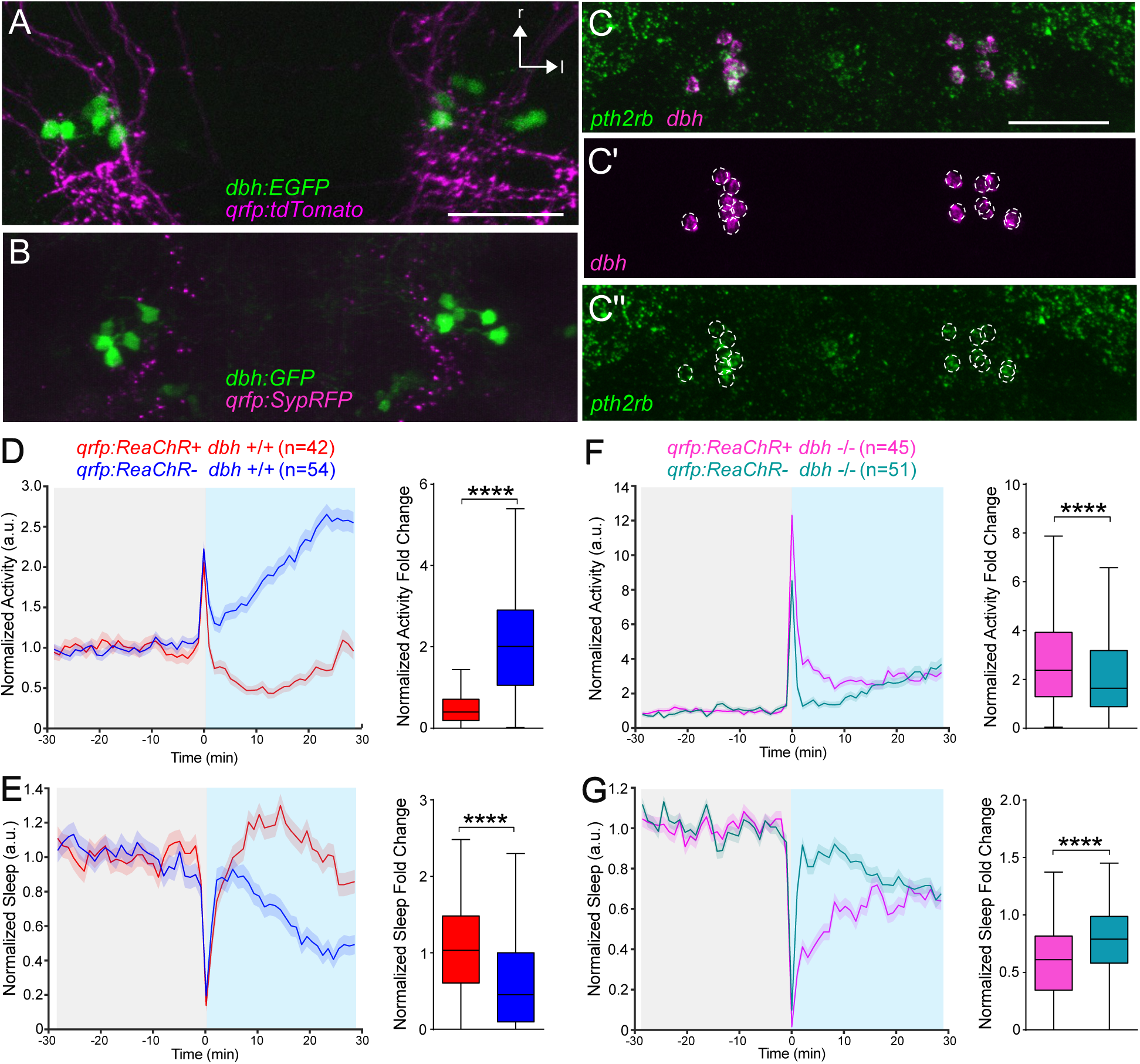
Sleep induced by optogenetic stimulation of Pth4 neurons is suppressed in *dbh* mutant fish. (**A-B**) Pth4 neuron fibers (**A**, tdTomato) and presynaptic terminals (**B**, synaptophysin-TagRFP (SypRFP)) are found in close proximity to the somata and neurites of GFP-labeled LC neurons. (**C-C”**) HCR for *dbh*, which labels LC neurons, and *pth2rb* shows that *pth2rb* is expressed in LC neurons (co-expression in 190/199 cells, n=18 fish). Scale bars: 50 μm. (**D-G**) Optogenetic stimulation of Pth4 neurons suppresses locomotor activity and increases sleep in *dbh* +/+ fish (**D,E**) but not in *dbh* -/- fish (**F,G**) compared to non-transgenic siblings. Mean ± SEM line plots and Tukey box plots are shown. n = number of fish. ****, p<0.0001 by Mann-Whitney U test.

**Figure 6.**
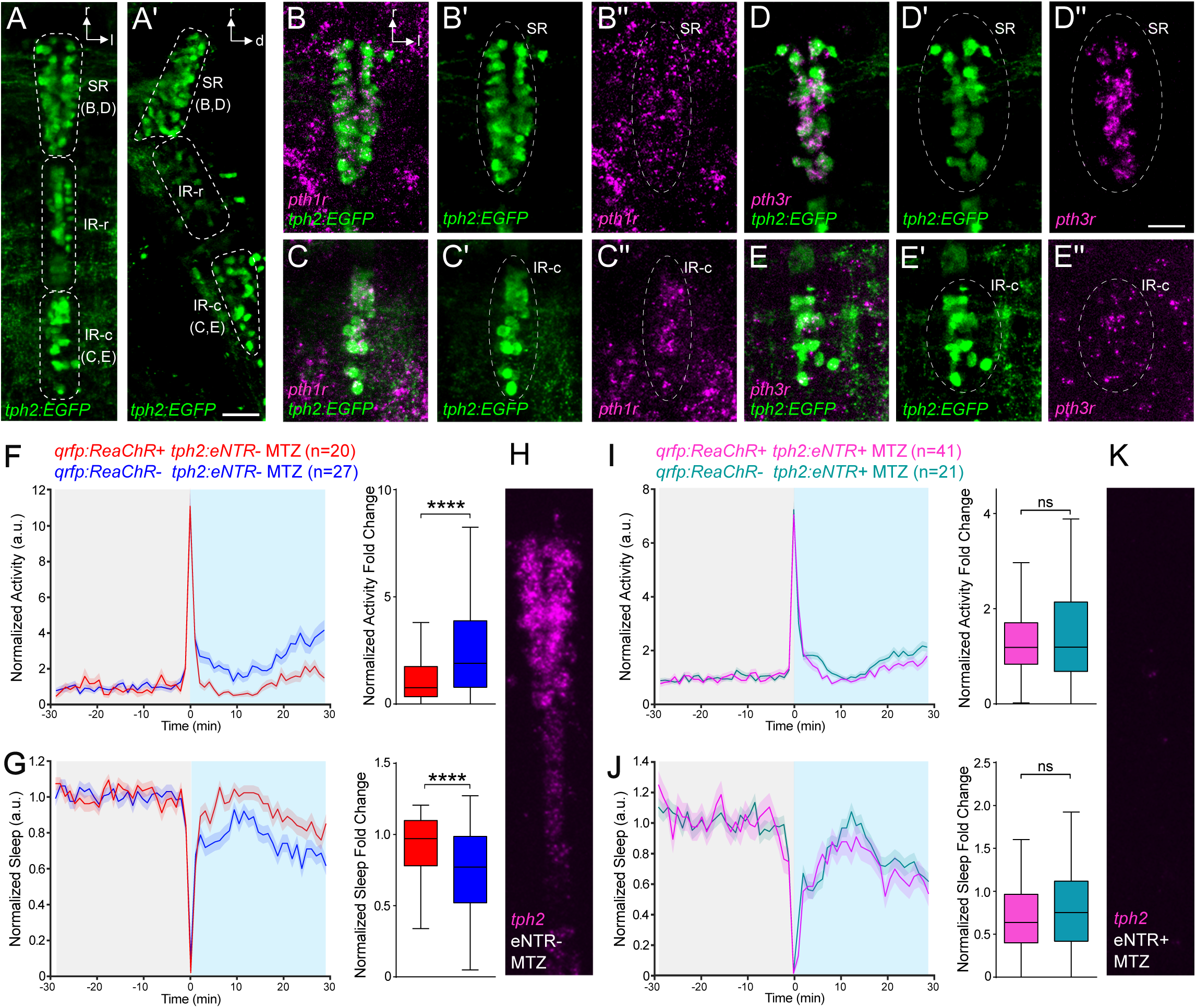
Sleep induced by optogenetic stimulation of Pth4 neurons is abolished by ablation of the 5HT raphe. (**A,A’**) 5HT raphe neurons labeled in *Tg(tph2:EGFP*) fish include the superior raphe (SR), rostral inferior raphe (IR-r) and caudal inferior raphe (IR-c) subregions, as seen in dorsal (**A**) and lateral (**A’)** views. (**B-E”**) HCR shows that *pth1r* (**B-C”**) and *pth3r* (**D-E”**) are both expressed in the SR (*pth1r*: in 97/226 SR neurons express *pth1r* in 5 fish, *pth3r*: 312/422 SR neurons express *pth3r* in 11 fish). Both receptors are also expressed in the IR-c (*pth1r*: 76/90 IR-c neurons express *pth1r* in 5 fish, *pth3r*: 59/82 IR-c neurons show faint expression of *pth3r* in 4 fish). Raphe neurons are indicated with dashed ovals. Scale bars: 25 μm. (**F-K**) Optogenetic stimulation of Pth4 neurons in *Tg(qrfp:ReaChR)* fish suppresses locomotor activity and increases sleep in MTZ-treated eNTR negative fish (**F,G**) with intact raphe (**H**), but not in MTZ-treated *Tg(tph2:eNTR)* fish (**I,J**) with ablated raphe (**K**), compared to ReaChR negative fish. Mean ± SEM line plots and Tukey box plots are shown. n = number of fish. ****, p<0.0001, ns, p>0.05 by Mann-Whitney U test. r, rostral; l, lateral; d, dorsal.

**Figure 7.**
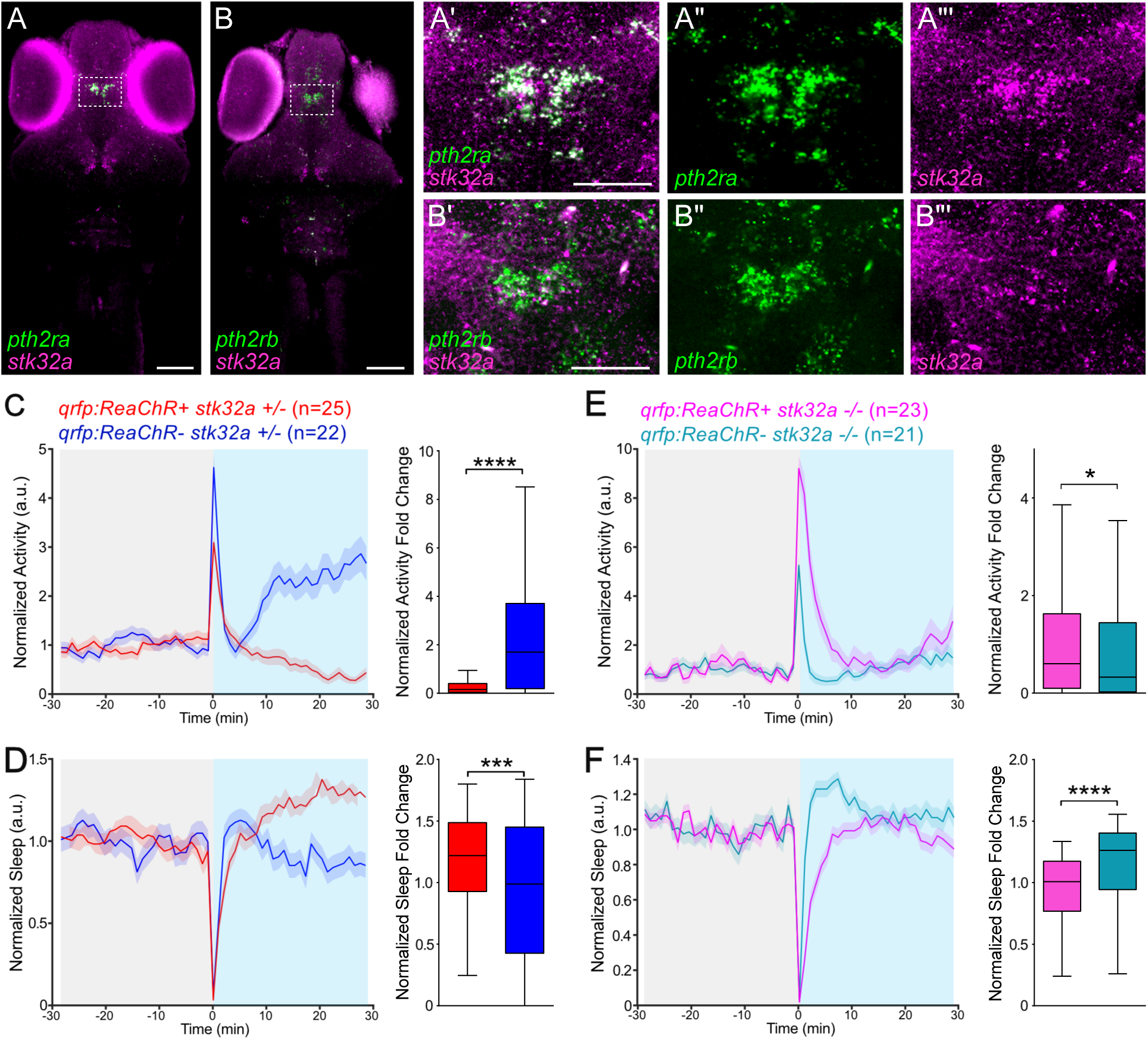
Sleep induced by optogenetic stimulation of Pth4 neurons is suppressed in *stk32a* mutant fish. (**A-B”**) HCR shows that *pth2ra* (**A-A”’**) and *pth2rb* (**B-B”’**) are both co-expressed with *stk32a* in the prethalamus. Boxed regions in (**A,B**) are magnified in (**A’-A”’**) and (**B’-B”’**) Scale bars: 100 μm (**A,B**); 50 μm (**A’,B’**). (**C-F**) Optogenetic stimulation of Pth4 neurons suppresses locomotor activity and increases sleep in *stk32a* +/- fish (**C,D**) but not in *stk32a* -/- fish (**E,F**) compared to ReaChR negative fish. Mean ± SEM line plots and Tukey box plots are shown. n = number of fish. *, p<0.05, ***, p<0.001, ****, p<0.0001 by Mann-Whitney U test.

### Pth4 neuron-induced sleep requires the LC

LC neurons are the main source of noradrenaline in the brain, and are essential for the wake-promoting function of Hcrt and Hcrt-expressing neurons in both mice ^51^ and zebrafish ^31^. Because Pth4 neurons project near the LC (Figures 1B, 1D and 5A), we explored a possible interaction between Pth4 and the LC in regulating sleep. First, we confirmed that *pthr2b*, but not *pth1r*, *pth2ra*, or *pth3r*, is expressed in the LC by performing HCR using *dopamine beta hydroxylase* (*dbh)*- specific probes to label LC neurons (Figures 5C-5C”, *pth2rb* is expressed in 190/199 LC neurons, n=18 fish; no co-expression of *dbh* with *pth1r*, *pth2ra*, or *pth3r*, Figure S3). Second, we generated transgenic fish in which Pth4 neurons express synaptophysin-TagRFP, which labels presynaptic terminals ^55^. We observed many synaptophysin-TagRFP puncta near the somata of LC neurons (Figure 5B), consistent with Pth4 neurons having synaptic contacts with LC neurons. Based on these observations, we hypothesized that Pth4 neurons promote sleep by inhibiting the function of LC neurons. If this hypothesis is correct, then the ability of Pth4 neurons to promote sleep should be suppressed in *dbh* mutants, which fail to synthesize noradrenaline and show increased sleep ^31^. We tested this prediction by comparing the effect of optogenetic stimulation of Pth4 neurons in animals that were either homozygous wild-type (WT) or homozygous *dbh* mutant. Consistent with our previous observations (Figure 2), stimulation of Pth4 neurons suppressed locomotor activity and increased sleep in *dbh* +/+ fish (Figures 5D and 5E). In contrast, these phenotypes were blocked, and even slightly reversed, when Pth4 neurons were stimulated in *dbh* -/- fish (Figures 5F and 5G). Stimulating Pth4 neurons in fish treated with the noradrenergic receptor antagonist prazosin gave similar results (data not shown). The apparently reversed nature of the phenotype may result from the hypersensitivity of *dbh* -/- fish to stimuli ^31^, such as the onset of blue light in this assay. These results are consistent with the hypothesis that Pth4 neurons promote sleep, in part, by inhibiting the function of LC neurons.

### Pth4 neuron-induced sleep requires the 5HT raphe

The 5HT raphe nuclei (RN) promote sleep homeostasis in both mice ^32^ and zebrafish ^32^, and are required for the sleep-promoting effect of NPVF neuron stimulation in zebrafish ^21^. We observed 3 subregions of the RN (Figures 6A and 6A’) with different *pth receptor* expression patterns. *pth1r* and *pth3r* (Figures 4A’, 4D’ and 6B-6E), but not *pth2ra* or *pth2rb* (Figure S4) are expressed in the superior raphe (SR, Figures 6B and 6D) and caudal inferior raphe (IR-c, Figures 6C and 6E). In contrast, the rostral inferior raphe (IR-r), does not express any *pth receptor*, but is innervated and activated by NPVF neurons ^21^. The expression of *pth1r* and *pth3r* in the RN suggests that Pth4 neurons may promote sleep via the RN. If this hypothesis is correct, then the ability of Pth4 neurons to promote sleep should be suppressed in fish whose RN are ablated. We tested this prediction by using *Tg(qrfp:ReaChR); Tg(tph2:eNTR)* fish to compare the effect of optogenetic stimulation of Pth4 neurons in animals that express enhanced nitroreductase (eNTR) ^56,57^ in the RN to their eNTR-negative siblings. We treated both populations of animals with 4 mM MTZ, which resulted in the complete loss of the RN in eNTR-positive animals (Figure 6K), but not in their eNTR-negative siblings (Figure 6H). Stimulation of Pth4 neurons suppressed locomotor activity and increased sleep in eNTR-negative animals whose RN are intact (Figures 6F and 6G), but these phenotypes were blocked in eNTR-positive animals whose RN were ablated (Figures 6I and 6J). These results are consistent with the hypothesis that Pth4 neurons promote sleep, in part, by promoting the sleep-inducing function of raphe neurons.

### Pth4 neuron-induced sleep requires *stk32a*

Based on human genetic data and a zebrafish genetic screen, mutation of *serine/threonine kinase 32a* (*stk32a*) was recently shown to increase sleep in both zebrafish and mice (Tran et al., submitted). This study also found that the neuropeptide *neurotensin* (*nts)* acts upstream of the 5HT RN in promoting homeostatic sleep pressure. We tested for interactions between *pth4* and *stk32a* in sleep for three reasons. First, the observations that the 5HT raphe is required for *pth4*- induced sleep (Figure 6) and *stk32a* is required for raphe-induced sleep (Tran et al., submitted) suggest that *stk32a* should also be required for *pth4*-induced sleep. Second, we found that *pth2ra* and *pth2rb* (Figures 7A and 7B), but not *pth1r* or *pth3r* (Figure S5), are co-expressed with *stk32a* in larval zebrafish prethalamic neurons, suggesting that Pth4 can signal via these neurons. Third, these prethalamic neurons also express *nts*, which promotes sleep by activating the 5HT raphe (Tran et al., submitted), and in a manner that requires both the 5HT raphe and *stk32a*. Based on these observations, we hypothesized that *stk32a* signaling is required for Pth4 neuron-induced sleep. We tested this hypothesis by comparing the effect of optogenetic stimulation of Pth4 neurons in animals that were either heterozygous or homozygous for a null mutation in *stk32a* (Tran et al., submitted). We found that the ability of Pth4 neuron stimulation to decrease locomotor activity and increase sleep in *stk32a* +/- fish (Figures 7C and 7D) was blocked, and slightly reversed, in *stk32a* -/- fish (Figures 7E and 7F). These results are consistent with the hypothesis that Pth4 neurons promote sleep, in part, via *stk32a*. They may do so either via prethalamic neurons that express *pth2ra, pth2rb*, and *stk32a*, or via other *stk32a*-expressing cells that act downstream of the 5HT raphe (Tran et al., submitted).

## Discussion

Here we identify neurons that express both *qrfp* and *pth4* as a hypothalamic neuronal population that promotes sleep in a manner that requires *pth4* but not *qrfp*. *pth4* belongs to the *pth* family of genes, which predate the evolution of vertebrates, as Pth-like peptides are also present in snail, cockroach, and amphioxus ^58^, and encode structurally-related secreted peptides that act via a family of G-protein coupled receptors ^52^. The mammalian genome contains three *Pth* genes, namely *Pth*, *Pth2/TIP39*, and *Pthlh*. The zebrafish genome has one or two paralogs for each of these mammalian genes, as well as *pth4*, an ancient *pth* family gene that is most similar to *pthlh*, and was lost in the placental mammalian lineage ^52^. Pth peptides regulate bone homeostasis and several developmental processes in vertebrates ^59–63^, and Pth2 has been shown to regulate social behaviors and anxiety in fish ^64,65^ and mice ^66,67^. Previous studies showed that both ablation of *pth4*-expressing neurons and overexpression of *pth4* in zebrafish resulted in abnormal bone mineralization ^41,42^, consistent with the expected function for a *pth* gene in regulating bone homeostasis. While PTH levels were shown to be temporally linked to sleep stage cycles in humans ^68^, to the best of our knowledge, this study is the first to show that a member of the Pth family of peptides can regulate sleep.

Consistent with an important role for Pth4 in regulating sleep is its exclusive expression in the hypothalamus, an ancient vertebrate brain region that is molecularly and anatomically conserved between zebrafish and mammals ^69–71^. A challenge of studying hypothalamic control of mammalian sleep is that in addition to sleep, the hypothalamus regulates several other important processes, including hunger, body temperature, circadian rhythms, and social behaviors ^6,72^. This complexity can make it difficult to distinguish direct effects of specific neuropeptides and neuronal populations on sleep from indirect effects via these other processes. In contrast, we use larval zebrafish before the onset of feeding ^73^, sexual differentiation ^74^ and complex behaviors such as social interactions ^75^, and thermoregulation is unlikely to be a factor in zebrafish sleep because they are poikilothermic. As a result, the use of larval zebrafish allows hypothalamic regulation of sleep to be studied without many of the complications that are inherent to mammalian models. Indeed, the use of larval zebrafish has uncovered direct roles for the hypothalamic neuropeptides Hcrt, NPY, MCH, and NPVF in regulating sleep ^13,19,20,30,31,43,76^. One caveat of this study is that we failed to detect an effect of mutation of *qrfp* and/or *pth4*, or ablation of *qrfp/pth4*-expressing neurons, on sleep. One possible interpretation of these results is that the effect of stimulating Pth4 neurons on sleep is a gain-of-function artifact that does not reflect the normal function of these neurons. However, there are several examples where gain-of-function perturbations produce robust sleep phenotypes, but corresponding loss-of-function perturbations produce small or no significant phenotype, in both zebrafish ^28,34,77^ and mice ^78,79^, likely due to compensatory or redundant mechanisms that regulate sleep.

We identified two mechanisms through which Pth4 neurons promote sleep. First, we found that stimulation of Pth4 neurons does not increase sleep in *dbh* mutants, which do not synthesize NA, or in animals treated with the NA receptor antagonist prazosin. Studies in zebrafish and rodents have shown that NA promotes arousal, and that inhibition of NA signaling increases sleep ^31,80^. Our observations thus suggest that Pth4 neurons promote sleep by inhibiting NA signaling. Consistent with this possibility, we found evidence for direct innervation of LC neurons by Pth4 neurons, and found that LC neurons express *pth2rb*. Similarly, a previous study found that NPY promotes zebrafish sleep by inhibiting NA signaling. NA signaling and the LC are also necessary for the wake-promoting effect of Hcrt neurons in both mammals ^51^ and zebrafish ^31^. Moreover, the neuropeptide neuromedin U promotes arousal via *corticotropin releasing hormone*-expressing neurons that are adjacent to the LC ^28^. Taken together, these observations suggest the LC, and possibly other nearby neuronal populations, integrate signals from multiple neuropeptidergic sleep- and wake-promoting neurons to determine vigilance state.

Second, we found that Pth4 neuron-induced sleep is abolished in fish whose 5HT raphe is ablated. This observation suggests that Pth4 neurons promote sleep by activating the raphe, whose stimulation promotes sleep in both zebrafish and mice ^32,81^, similar to stimulation of Pth4 neurons in zebrafish. Consistent with this hypothesis, we found that *pth1r* and *pth3r* are expressed in superior raphe and caudal inferior raphe neurons. We previously showed that NPVF neurons, whose somata are adjacent to those of Pth4 neurons in the hypothalamus, promote sleep by activating rostral inferior raphe neurons. Our observation that mutation of *npvf* does not suppress the sedating effect of stimulating Pth4 neurons (data not shown) suggests that NPVF and Pth4 neurons independently promote sleep by stimulating different subpopulations of 5HT raphe neurons. These results suggest that the 5HT raphe integrates signals from multiple neuropeptidergic sleep-promoting neurons to determine vigilance state, similar to the LC. We recently found that loss of the protein kinase *stk32a* results in increased sleep, and that *stk32a* regulates sleep homeostasis in the same genetic pathway as the 5HT raphe (Tran et al., submitted). We also showed that neurons in the prethalamus co-express the neuropeptide *nts* and *stk32a*, and that activation of Nts signaling promotes sleep by stimulating neurons in the superior raphe. Our findings in this study that stimulation of Pth4 neurons does not enhance the increased sleep observed in *stk32a* mutants, and that the prethalamic neurons that co-express *stk32a* and *nts* also express *prh2ra* and *pth2rb*, suggest that Pth4 neurons may promote sleep by activating prethalamic *stk32a/nts* neurons, which results in activation of the 5HT raphe, leading to increased sleep. Alternatively, or in addition to this mechanism, Pth4 neurons may promote sleep via direct interaction with the 5HT raphe since both *pth1r* and *pth3r* are expressed in superior raphe and caudal inferior raphe neurons. Our proposed sleep circuitry model is illustrated in Figure 8.

**Figure 8.**
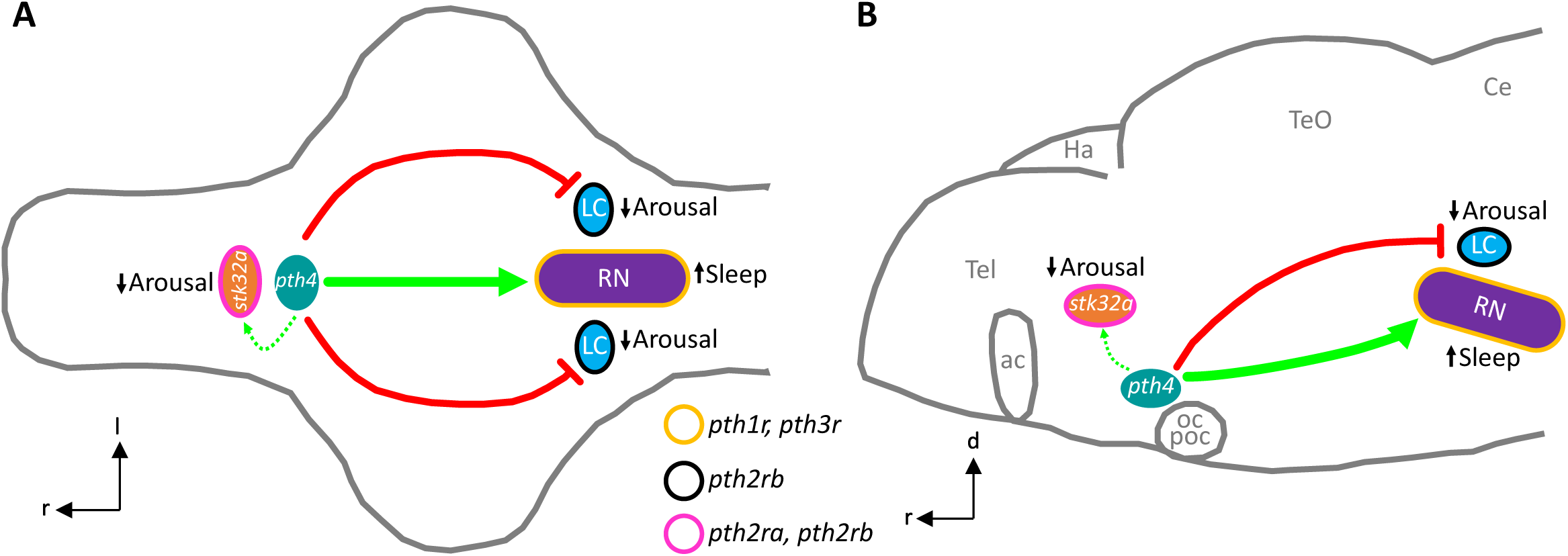
Model for sleep regulation by Pth4 neurons. Pth4 neurons in the hypothalamus promote sleep by activating the sleep-promoting 5HT SR and inhibiting the arousal-promoting LC in the hindbrain. Pth4 neurons also functionally interact with *stk32a*, possibly by activating neurons in the prethalamus that express both *stk32a* and *nts*, and/or via other *stk32a*-expressing cells. Colored outlines of neuronal populations indicate expression of specific *pth* receptors. Green arrows indicate activation. Dashed green arrows indicate potential activation. Red lines indicate inhibition. Dorsal (**A)** and lateral (**B**) views are shown. Gray lines show the outline of the brain and anatomical landmarks. ac, anterior commissure; Ce, cerebellum; Ha, habenula; LC, locus coeruleus; oc, optic chiasm; poc, postoptic commissure; RN, raphe nuclei; Tel, telencephalon; TeO, optic tectum. l, lateral; d, dorsal; r, rostral.

While we identified post-synaptic mechanisms through which Pth4 neurons promote sleep, the mechanisms that regulate Pth4 neurons remain unknown. Pth4 neurons have extensive short fibers throughout the hypothalamus, which contains several neuronal populations that promote either arousal (e.g. hypocretin ^31^, neuromedin U ^28^) or sleep (e.g. NPVF ^20,21^, MCH ^19^, prokineticin 2 ^82^), and may play a role in regulating Pth4 neurons. The presence of Pth4 neuron fibers in the optic tectum, which processes visual information and is homologous to the mammalian superior colliculus, and the ventral pretectum, which also receives retinal input and responds to lights off stimuli ^83^, suggest that Pth4 neurons might play a role in the effects of light and dark on sleep. Future experiments that explore mechanisms that regulate the activity of Pth4 neurons are needed to address these questions.

Mammals lack a Pth4 ortholog, so the relevance of our zebrafish findings to mammalian sleep is unclear. However, a recent study found that stimulation of *qrfp*-expressing neurons induces a torpor-like state in mice, in which locomotor activity, body temperature, and oxygen consumption are dramatically reduced ^84^. Together, the zebrafish and rodent data suggest that QRFP neurons serve an evolutionarily conserved role in energy conservation, which takes the form of sleep in poikilothermic fish, which do not regulate their body temperature, and a torpor-like state in homeothermic rodents. Notably, the QRFP peptide is not required for the induction of sleep in zebrafish or torpor in mice. Instead, we found that *pth4* is required for QRFP/Pth4 neuron-induced sleep in zebrafish, while both glutamate and GABA are important for QRFP neuron-induced torpor in mice. An interesting difference between zebrafish and mice concerns the location of QRFP neurons in the hypothalamus. In zebrafish, QRFP/Pth4 neurons comprise a small cluster of hypothalamic neurons that are intermingled with Hcrt neurons, in a brain region that corresponds to the mammalian lateral hypothalamus (LHA). In mice, *qrfp* is expressed not only in the LHA, but also in the anteroventral periventricular nucleus, medial preoptic area, and paraventricular nucleus of the hypothalamus, and it is stimulation of these non-LHA populations that can induce torpor, whereas stimulation of QRFP neurons in the LHA does not ^84^. These observations suggest that expression of QRFP within the hypothalamus has expanded in the mammalian lineage, and the new expression domains in mammals may have enabled in a shift in the function of QRFP neurons from promoting sleep to inducing torpor. Another potential difference between zebrafish and mice is that *hcrt* and *qrfp* are not co-expressed in zebrafish, but two studies observed co-expression of the two genes in the mouse LHA ^85,86^, although a third study found that *hcrt* and *qrfp* comprise distinct populations in the mouse LHA ^87^. Furthermore, since the QRFP peptide does not promote sleep in either zebrafish or mice ^34,47^, the relevance of any *hcrt* and *qrfp* co-expression to sleep is unclear, and the function of QRFP that might be expressed in mouse Hcrt neurons is unknown.

Our discovery that Pth4 neurons promote sleep is interesting in light of previous studies demonstrating a role for these neurons in bone physiology, particularly in bone mineralization and phosphate homeostasis. Earlier studies in zebrafish showed that ablation of *pth4*-expressing neurons leads to reduced bone mineralization ^41,42^, while overexpression of *pth4* results in decreased bone mineral density ^41^. Interestingly, although mammals lost the *pth4* gene during evolution, likely due to functional redundancy with *pth1*, similar roles in bone biology have been observed. For example, human PTH increases bone resorption and the healing of bone fractures in mice ^88^, and is used to treat or prevent osteoporosis in humans ^89,90^. A potential connection between sleep and bone regulation is supported by mammalian studies. For example, sleep restriction decreases bone mineral density in rats ^91,92^ and bone formation in humans ^93–95^, and both increased and decreased sleep are associated with an increased risk of osteoporosis in humans ^96–99^. Taken together, our study and these published studies suggest that Pth4 might link sleep and bone homeostasis.

In summary, we have identified a population of hypothalamic neurons that express the neuropeptide *pth4* whose stimulation results in a large increase in sleep, and showed that these neurons promote sleep via interactions with monoaminergic populations in the brainstem. Additional studies are required to determine whether any of the mammalian Pth neuropeptides, and the neurons that express them, play roles in regulating mammalian sleep. Studies of the arousing neuropeptide Hcrt have led to the development of small molecule Hcrt receptor antagonists that are effective treatments for insomnia ^100^. If the role of zebrafish *pth4* in sleep is conserved for a member of the mammalian family of *Pth* genes, they could similarly serve as a target for novel therapies to treat sleep disorders.

## METHODS

### Key resources table

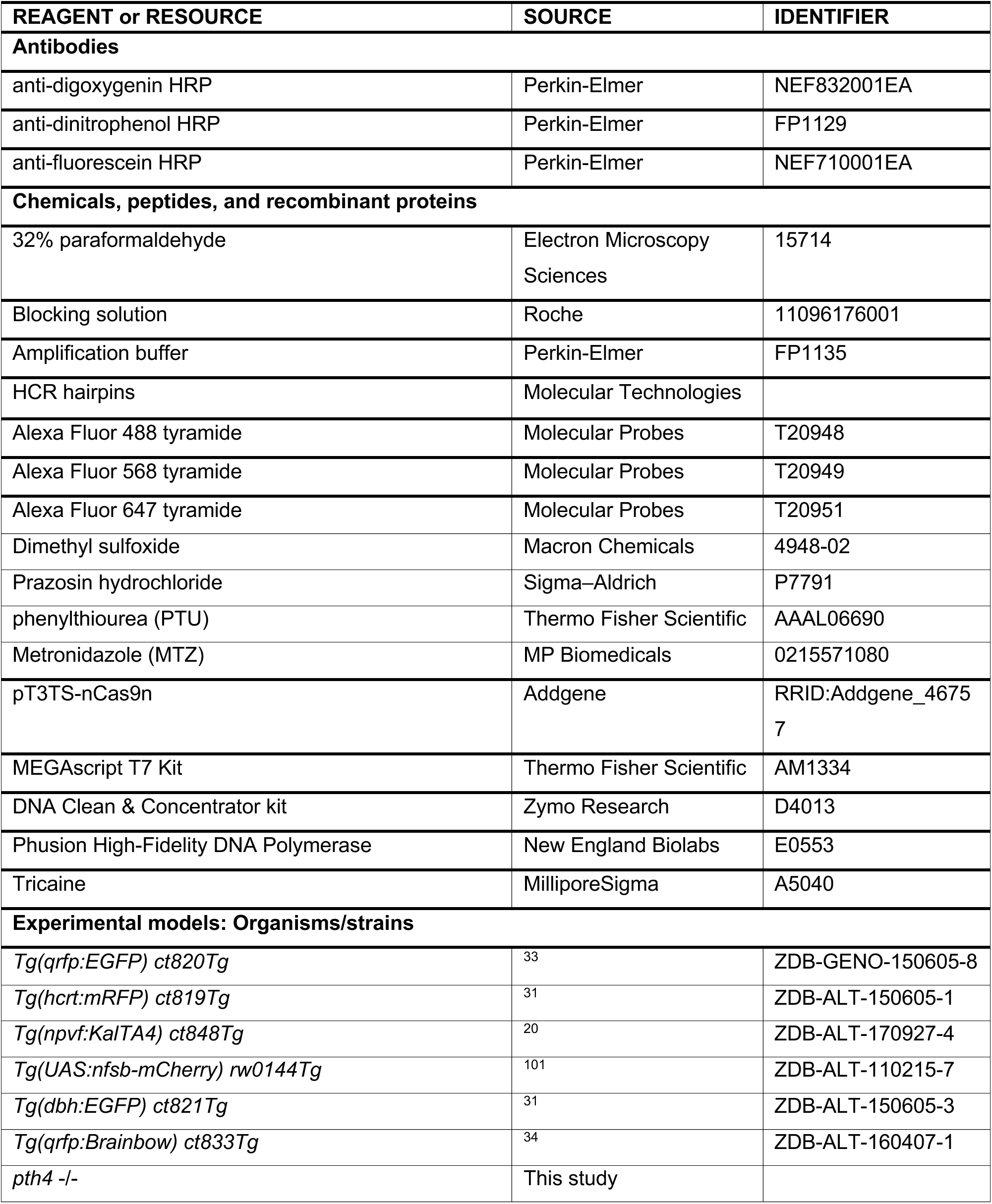
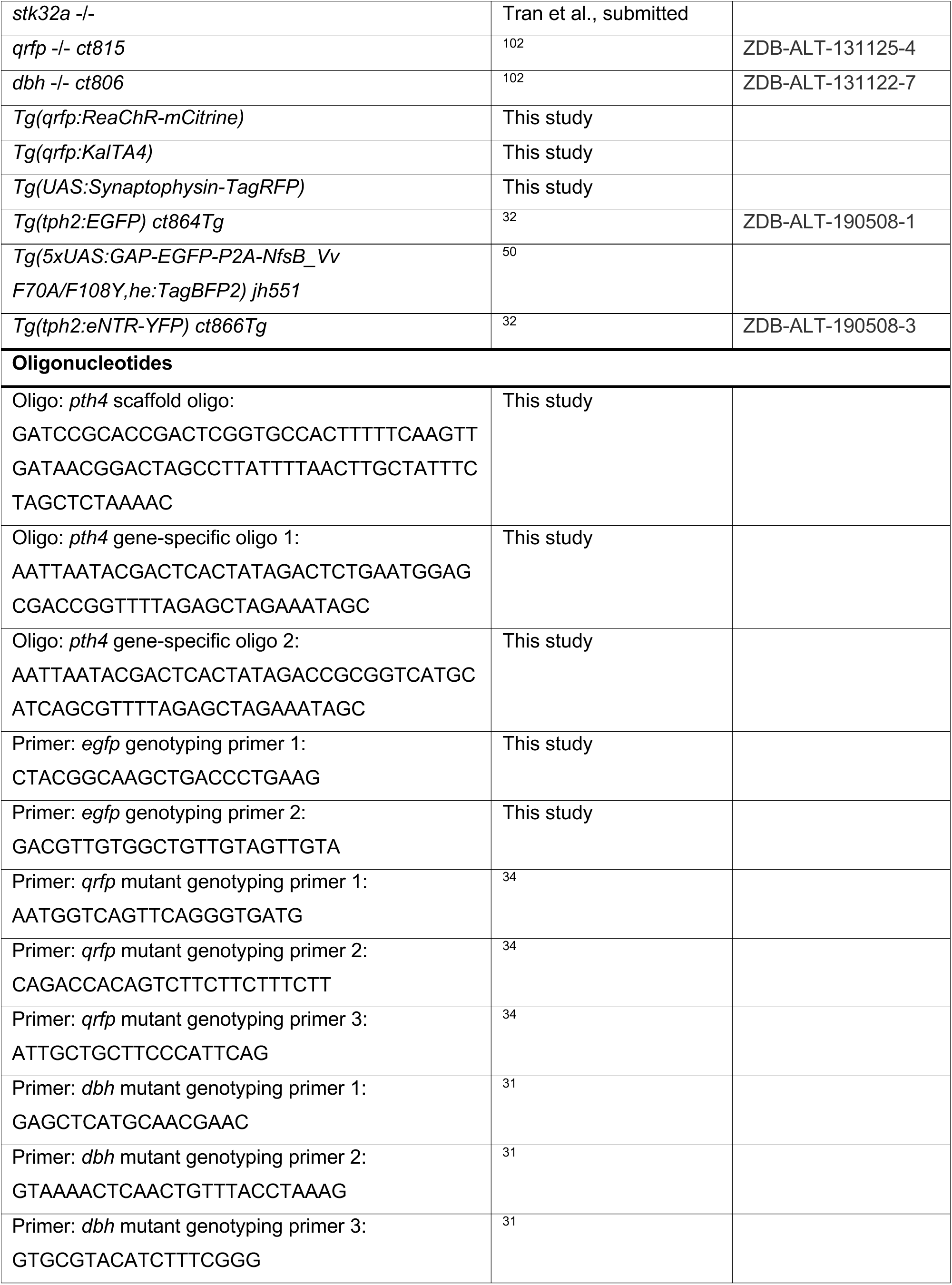
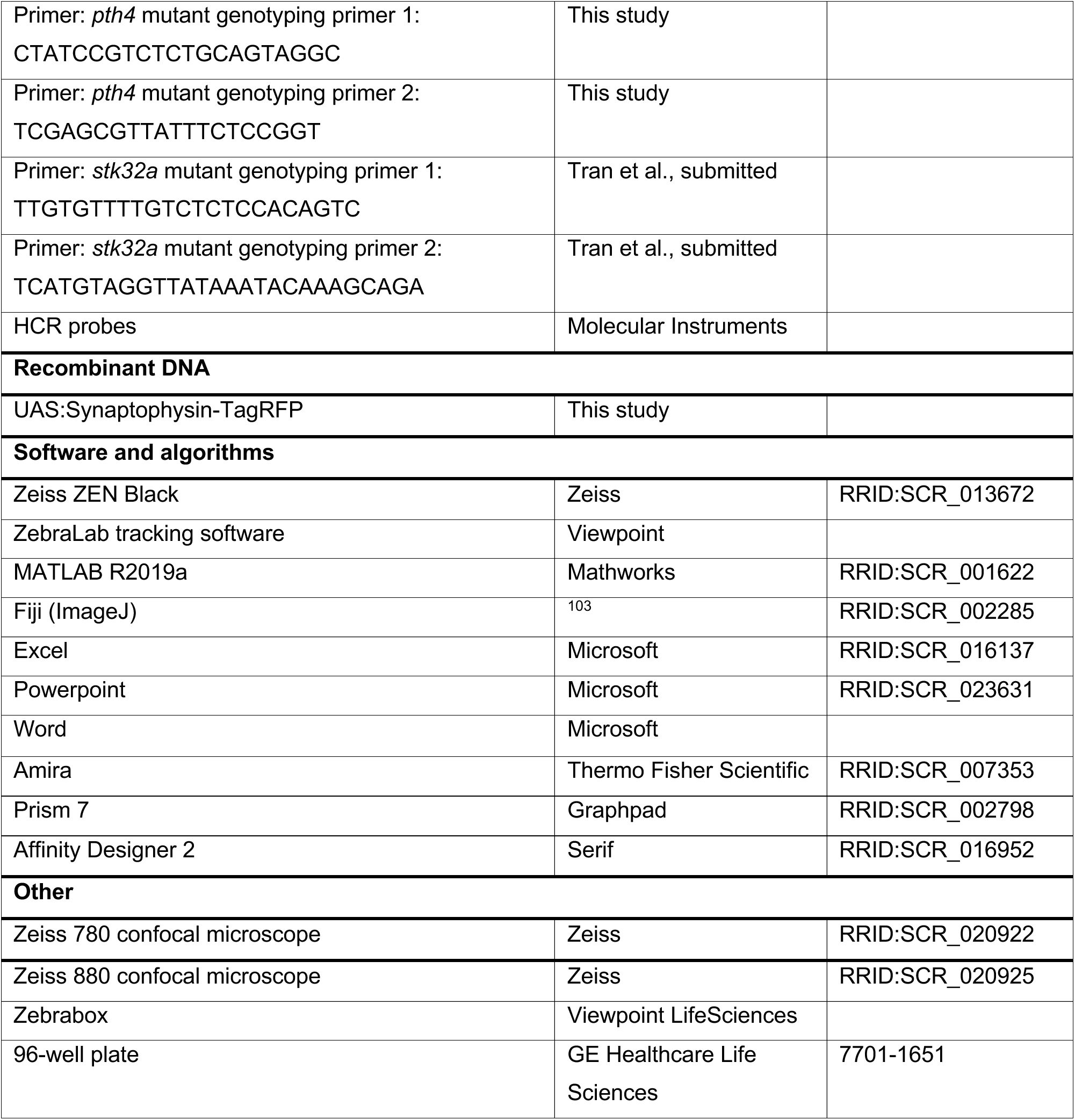

## EXPERIMENTAL MODEL AND SUBJECT DETAILS

### Zebrafish husbandry

All zebrafish experiments were performed in accordance with the Institutional Animal Care and Use Committee (IACUC) guidelines and by the Office of Laboratory Animal Resources at the California Institute of Technology (animal protocol 1836). Zebrafish from the TLAB strain were generated by mating AB and TL strains obtained from the Zebrafish International Resource Center (ZIRC, Oregon). TLAB (wild-type) animals were used to generate mutants and transgenic animals. Adult zebrafish were maintained on a 14:10 hour light:dark cycle with lights on at 9 a.m. and off at 11 p.m. All zebrafish experiments used healthy and experimentally naïve larval zebrafish at 5-6 dpf, a developmental stage prior to sexual differentiation. Zebrafish larvae were raised in petri dishes with 50 animals per plate containing E3 medium (5 mM NaCl, 0.16 mM KCl, 0.33 mM CaCl2, and 0.33 mM MgSO4).

### Zebrafish mutagenesis

The *pth4* mutant was generated using CRISPR/Cas9 by co-injecting two sgRNAs targeting exon 2 of the *pth4* gene (designed with CHOPCHOP v2 ^104^) and *cas9* mRNA (50 ng/µL, transcribed from the pT3TS-nCas9n plasmid, Addgene) into WT zebrafish embryos at the one-cell stage. sgRNAs were synthesized via template-free PCR using a universal scaffold oligo and gene-specific oligos, followed by *in vitro* transcription with the MEGAscript T7 Kit (Ambion) and purification with the Zymo DNA Clean & Concentrator-5 Kit (Zymo Research). The PCR used the following oligonucleotides (IDT): scaffold oligo: 5′- GATCCGCACCGACTCGGTGCCACTTTTTCAAGTTGATAACGGACTAGCCTTATTTTAACTTG CTATTTCTAGCTCTAAAAC-3′; gene-specific oligo 1 (GS1): 5’ - AATTAATACGACTCACTATAGACTCTGAATGGAGCGACCGGTTTTAGAGCTAGAAATAGC-3′; gene-specific oligo 2 (GS2): 5’ - AATTAATACGACTCACTATAGACCGCGGTCATGCATCAGCGTTTTAGAGCTAGAAATAGC-3′. Approximately 2 nL of the injection mix, containing 25 ng/µL of each sgRNA, *cas9* mRNA, and 0.1% phenol red, was microinjected into one-cell stage embryos using a pressure injector (ASI MPPI-2). Uninjected embryos served as controls. Sanger sequencing revealed a 17 bp deletion that caused a frameshift and introduced a premature stop codon, resulting in a truncated 53 amino acid protein that lacks the Pth-related domain and is likely a complete loss-of-function allele. A stable line carrying this mutation, *pth4 iim21*, was established and used for all experiments. The *qrfp^ct815^* ^34^ and *dbh^ct806^* ^31,102^ mutants have previously been described. The *stk32a* mutant is described in a manuscript that is under review (Tran et al., submitted).

### Zebrafish transgenesis

We generated *Tg(qrfp:ReaChR-mCitrine)* and *Tg(qrfp:KalTA4)* transgenic fish by cloning a 1 kb *qrfp* promoter ^33^, and ReaChR ^49^ fused to mCitrine ^105^ or KalTA4 ^106^, and SV40-polyA into a plasmid containing NRSE elements ^107,108^ and flanking Tol2 transposase recognition sequences ^109^. We generated *Tg(UAS:synaptophysin-TagRFP)* transgenic fish by cloning a *UAS*:*synaptophysin-TagRFP* transgene into a plasmid containing Tol2 transposase recognition sequences. We generated stable transgenic lines by co-injecting the plasmids with *tol2 transposase* mRNA into embryos at the single cell stage. Transgenic animals were identified by fluorescent tag expression or PCR. We used the following published lines: *Tg(qrfp:EGFP)ct820* ^33^, *Tg(hcrt:mRFP)ct819* ^33^, *Tg(npvf:KalTA4)ct848* ^20^, *Tg(UAS:nfsb-mCherry)rw0144* ^101^, *Tg(dbh:EGFP)ct821* ^33^, *Tg(qrfp:Brainbow)ct833* (used to label QRFP/Pth4 neurons with tdTomato) ^34^, *Tg(tph2:EGFP)ct864* ^32^, *Tg(tph2:eNTR-mYFP)ct866* ^32^, *Tg(5xUAS:GAP-EGFP-P2A-NfsB_Vv F70A/F108Y,he:TagBFP2) jh551* ^50^.

## METHOD DETAILS

### Pharmacology

Prazosin hydrochloride was dissolved in dimethyl sulfoxide (DMSO) and added to E3 medium at a concentration of 100 μM prazosin and 0.1% DMSO. Controls were treated with 0.1% DMSO in E3 medium.

### Sleep behavioral experiments

Sleep behavioral experiments were performed using an automated videotracking system as previously described ^13^. Zebrafish larvae were raised in petri dishes (max. 50 per dish) on a 14:10 hour light:dark (LD) cycle at 28.5°C with lights on a 9 a.m. and off at 11 p.m. Starting at 4 dpf, individual larvae were placed in each well of a 96-well plate (7701-1651, Whatman), filled with 650 µL E3 medium. Locomotor activity was quantified using an automated videotracking system (Viewpoint Life Sciences) with a Dinion one-third Monochrome camera (Dragonfly 2, Point Grey) fitted with a variable-focus megapixel lens (M5018-MP, Computar) and an infrared filter. The movement of each fish was recorded using quantization mode. The 96-well plate and camera were housed inside a custom Zebrabox (Viewpoint Life Sciences), with white light illumination from 9 a.m. to 11 p.m., and continuous infrared light illumination for video recording. The 96-well plate was housed in a chamber filled with recirculating water to maintain a constant temperature of 28.5°C. The parameters used for detection (detection threshold: 15, burst: 29, freeze: 3, bin size: 60 s) were empirically determined. Behavior was monitored for 24 hours, during the 6th day and night of development. Data were analyzed using custom MATLAB (Mathworks) scripts ^110^. Zebrafish were genotyped by PCR after each experiment to identify mutant, transgenic, and WT siblings.

### Optogenetic stimulation

The videotracking system was modified to include an array of blue LEDs (470 nm, MR-BO0040-10S, Luxeon V-star) mounted 15 cm above and 7 cm away from the center of the 96-well plate to ensure uniform illumination, as previously described ^31^. The LEDs were controlled using a custom Arduino microcontroller. A power meter (1098293, Laser-check) was used before each experiment to confirm light intensity (1.2 mW/cm²). In the afternoon at 4 dpf, zebrafish were placed in each well of a 96-well plate in a videotracker, and allowed to habituate overnight with exposure to 30 minute blue light stimuli at 12 a.m. and 12 p.m. On the 6th night of development, 6 trials were performed in which fish were exposed to blue light for 30 minutes, with 60 minutes of recovery between each trial, starting at 12 a.m. In some experiments trials continued throughout the day until the end of the experiment. The behavior of each fish was monitored for 30 minutes before and after blue light onset. Since the onset and offset of light stimuli induce startle responses with a short burst of locomotor activity, we excluded 3 minutes of data after light onset, and 2 minutes of data before light offset, from the analysis. For the baseline, we used a time period equal to blue light exposure, but prior to blue light onset.

### Chemogenetic ablation

Fish were treated with 4 mM metronidazole (MTZ; MP Biomedicals, 0215571080) diluted in E3 medium containing 0.2% DMSO from 2-4 dpf, refreshed every 24 hours, as previously described ^32^. Fish were kept in dim red light during the day (by using white light illumination and wrapping the plate in a red filter) throughout the treatment period to prevent MTZ photodegradation. At 4 dpf, fish were rinsed three times with E3 medium and allowed to recover for at least 1 hour prior to being transferred to 96-well plates for VT experiments. The reported data is from the 6th day and night (sleep/wake assay) or the 6th night and 7th day (optogenetic assay) of development.

### Fluorescence in situ hybridization (FISH) and hybridization chain reaction (HCR)

FISH ^111^ and HCR ^112^ were performed as previously described. Whole zebrafish larvae were fixed in 4% PFA in PBS overnight with nutation at 4°C, and subsequently washed with 0.1% Tween-20/PBS (PBSTw). Samples were permeabilized with graded dilutions of 100% methanol at −20°C for 10 minutes, rehydrated with graded dilutions of methanol/PBSTw, and washed with 0.25% Triton X-100/PBS (PBSTx). To improve transparency, samples were treated with a bleaching solution (3% hydrogen peroxide and 0.5% potassium hydroxide) for 20 minutes, and then washed with PBSTx. Samples were pre-hybridized for 60 minutes and then hybridized with riboprobes (FISH) or HCR probes (HCR, Molecular Technologies, 2 pmol of each probe set in 500 μL) overnight at 68°C (FISH) or at 37°C for 72 hours (HCR). Samples were washed with a wash buffer (Molecular Technologies) at 37°C (HCR) and equilibrated with 2x and then 0.2x (FISH) or 5x (HCR) SSCTx. Samples were then transferred to a blocking buffer (Roche) and incubated with HRP antibodies at 4°C (FISH), or to an amplification buffer (Molecular Technologies) with 30 pmol of snap-cooled hairpins at room temperature (HCR) in the dark overnight. For both FISH and HCR (for receptors that were expressed at low levels), tyramide signal amplification was performed using HRP antibodies and tyramides. HCR tyramide signal amplification uses standard HCR probes with hairpins that have binding sites for peroxidase-conjugated antibodies as used for FISH (Singh et al, manuscript in preparation). Samples were then washed with PBSTx, and gradually transferred to 80% glycerol in PBS prior to imaging.

### Confocal microscopy

Live animals were embedded in 1.5% low-melt agarose, while fixed animals were mounted in 80% glycerol. 3D stacks were recorded using Zeiss 780 and 880 confocal microscopes. Each channel was recorded sequentially, using alternating excitation wavelengths specific for each fluorophore, to avoid interfering signals from overlapping emission spectra. Zoom, dimensions, gain, offset, average, and speed were adjusted for each stack to obtain the optimal image quality of the desired volume. Stacks were evaluated using Fiji to create maximum intensity projections of the volume of interest, excluding signals from planes above or below. Brightness and contrast were adjusted for each channel. For some samples in which skin autofluorescence obscured the underlying neuronal signal of interest, Amira (Thermo Fisher Scientific) was used to digitally remove the skin using the Volume Edit module. Rotation animations were performed using the Movie Maker module. Amira was used in the Beckman Institute Resource Center for Transmission Electron Microscopy at Caltech.

## QUANTIFICATION AND STATISTICAL ANALYSIS

For all behavioral experiments the unit of analysis for statistics is a single animal, except for optogenetic experiments where the unit of analysis for statistics is a single trial. Line graphs represent mean ± SEM. Tukey box plots are used for data presentation and data points outside the Tukey range are not shown to facilitate data presentation, but were included in statistical analyses. In optogenetic experiments, outlier data points outside 4x the inter-quartile-range were excluded from the analysis. Statistical analyses were performed using Prism 7 (GraphPad). Shapiro-Wilk normality tests indicated that behavioral data were not normally distributed. Therefore, we used non-parametric tests for statistical analyses (Mann-Whitney U test for 2 unpaired groups, Kruskal-Wallis test with Dunn’s correction for multiple comparisons for more than two unpaired groups). Data are considered to be statistically significant if p<0.05.

## ACKNOWLEDGMENTS

We thank Tasha Cammidge, Uyen Pham, Hannah Hurley, Brianna Garcia, Cristina Gonzalez, and Viveca Sapin for technical assistance; Alex Mack, Chris Cook, Caressa Wong, Barbara Orozco, Axel Dominguez, and Daisy Chilin for zebrafish care; Alex Schier and Constance Richter for sharing unpublished *pth4* mutants; and all members of the Prober Lab for comments and suggestions. This work was supported by grants from the German Research Foundation (U.H. DFG HE 7815/1-1), the Max Planck Society (S.R.), the Spanish Ministry of Science and Innovation (J.R. MCIN/AEI/10.13039/501100011033, AGL2017-89648P, and “ERDF A way of making Europe”), and the NIH (D.A.P. R35 NS122172).

## AUTHOR CONTRIBUTIONS

Conceptualization: UH, DAP

Methodology: UH, ST, CS, GO, SR, JR

Investigation: UH

Visualization: UH, DAP

Funding acquisition: SR, JR, DAP

Project administration: DAP

Supervision: DAP

Writing – original draft: UH, DAP

Writing – edit and revision: UH, SR, JP, DAP

## SUPPLEMENTAL FIGURE LEGENDS

**Figure S1.**
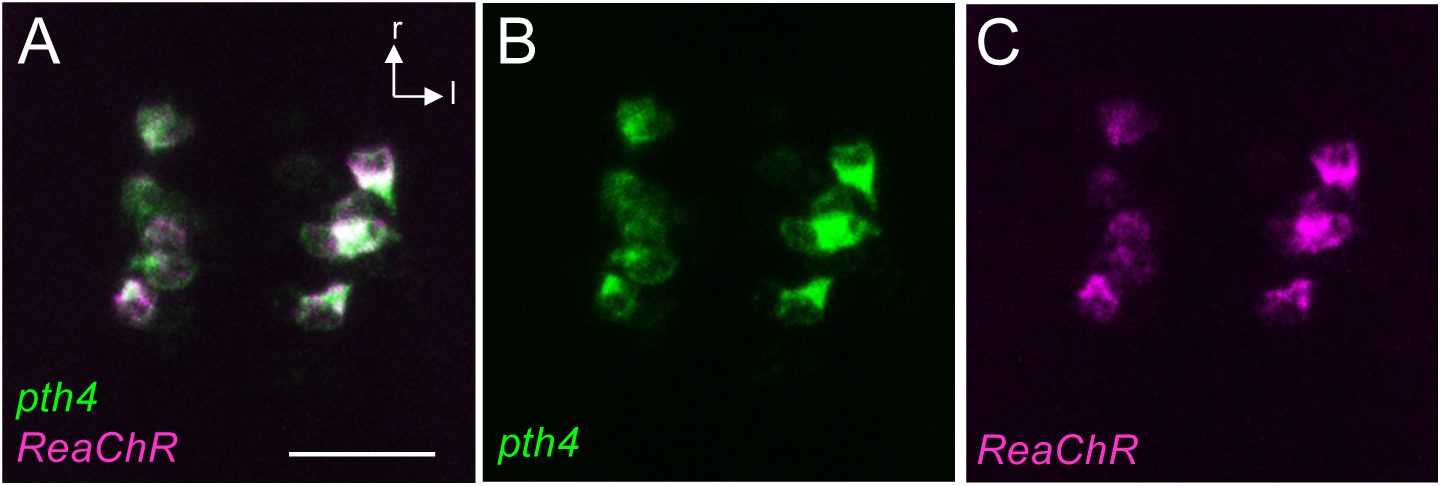
The *qrfp:ReaChR-mCitrine* transgene is expressed in QRFP/PTH4 neurons. HCR using probes specific for *pth4* (green, **A,B**) and *mCitrine* (magenta, **A,C**) shows that *ReaChR-mCitrine* is specifically expressed in QRFP/Pth4-expressing neurons in *Tg(qrfp:ReaChR-mCitrine)* fish (247/296 of Pth4 neurons express *ReaChR-mCitrine* in 13 fish). Scale bar: 25 μm. r, rostral; l, lateral.

**Fig S2.**
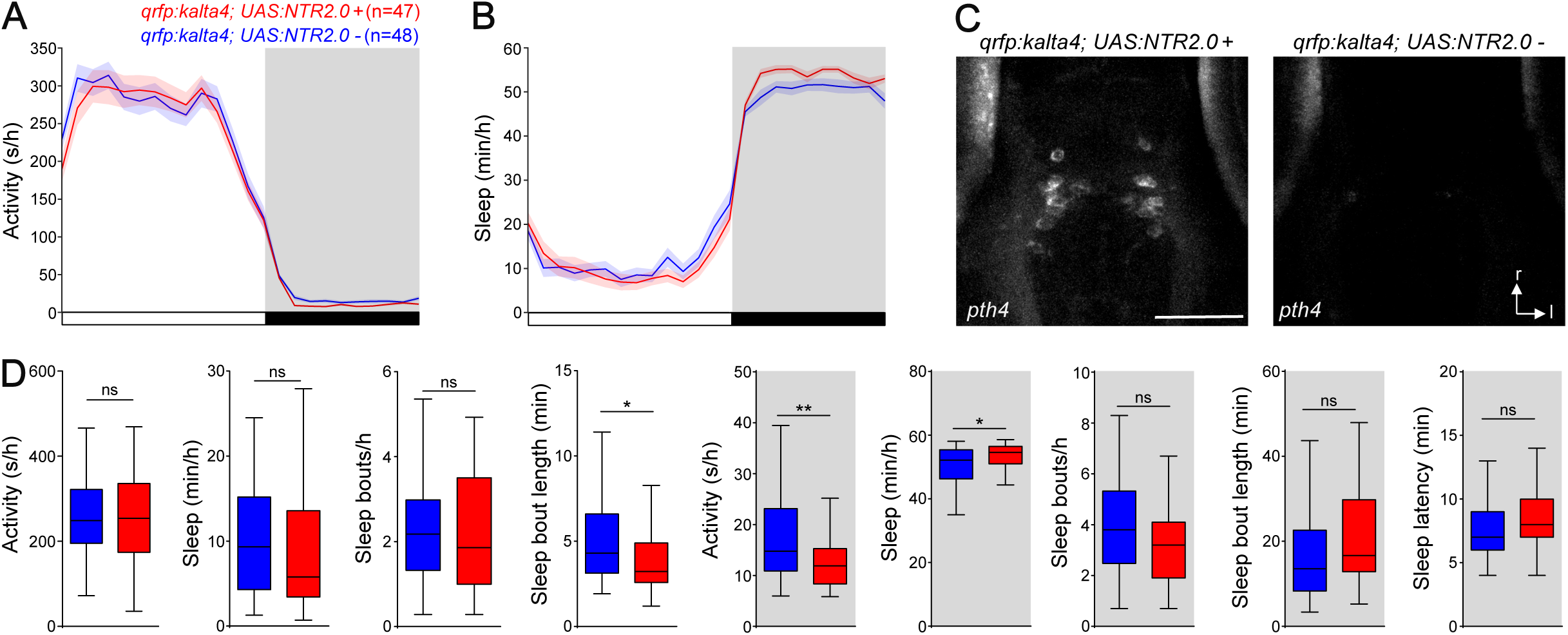
Locomotor activity and sleep are unaffected by *pth4* mutation or Pth4 neuron ablation. Ablation of *pth4*-expressing cells due to expression of NTR 2.0 and treatment with 4 mM MTZ results in a small decrease in locomotor activity (**A, D**) and increase in sleep (**B, D**) at night compared to identically treated non-transgenic siblings. Most other sleep parameters were unaffected by ablation of *pth4*-expressing cells (**D**). Ablation of *pth4*-expressing cells was confirmed by performing HCR using *pth4*-specific probes (**C**). Scale bar: 50 μm. r, rostral; l, lateral. n = number of fish. *, p<0.05, **, p<0.01 by Mann-Whitney U test.

**Figure S3.**
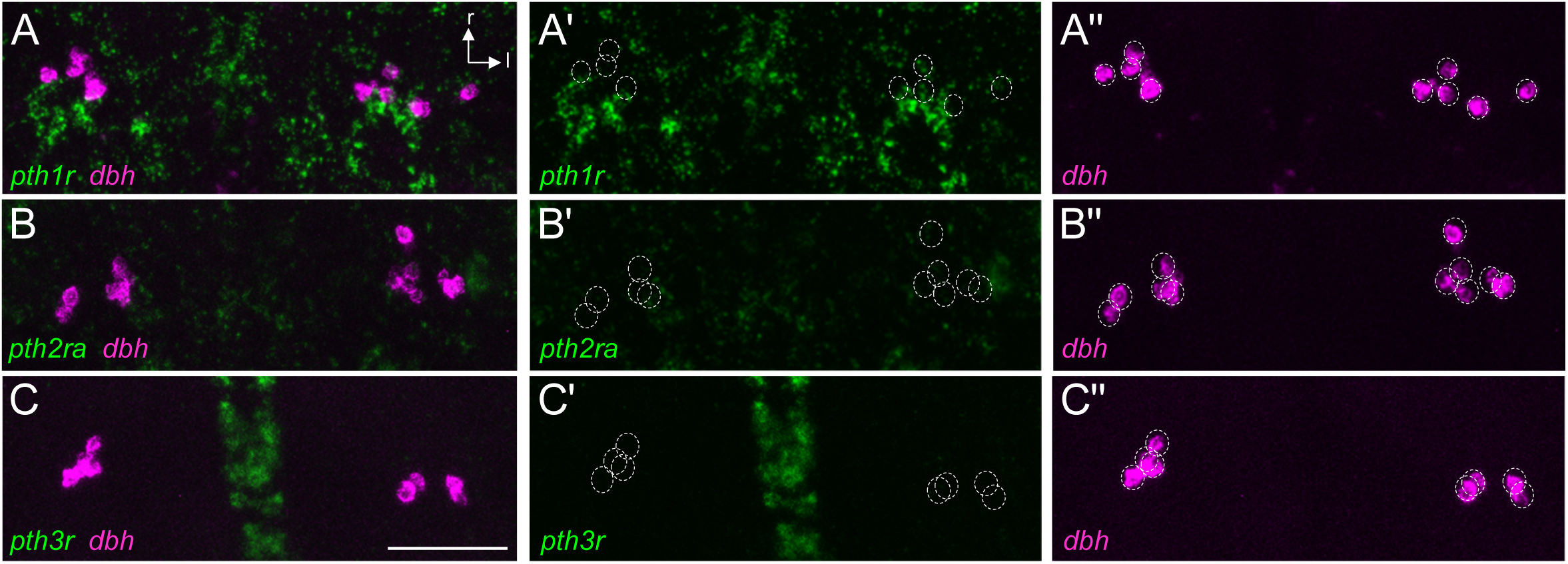
*pth1r*, *pth2ra*, and *pth3r* are not expressed in LC neurons. HCR using probes specific for *pth1r* (**A,A’**), *pth2ra* (**B,B’**), *pth3r* (**C,C’**), and *dbh* (**A,A”,B,B”,C,C”**) shows that none of these *pth receptors* is co-expressed with *dbh*, which labels LC neurons. *pth1r*: 0/61 *dbh* neurons express *pth1r* in 5 fish. *pth2ra*: 0/65 *dbh* neurons express *pth2ra* in 6 fish. *pth3r*: 0/88 *dbh* neurons express *pth3r* in 8 fish. Dashed circles indicate locations of *dbh*-expressing LC neurons. Scale bar: 50 μm. r, rostral; l, lateral.

**Figure S4.**
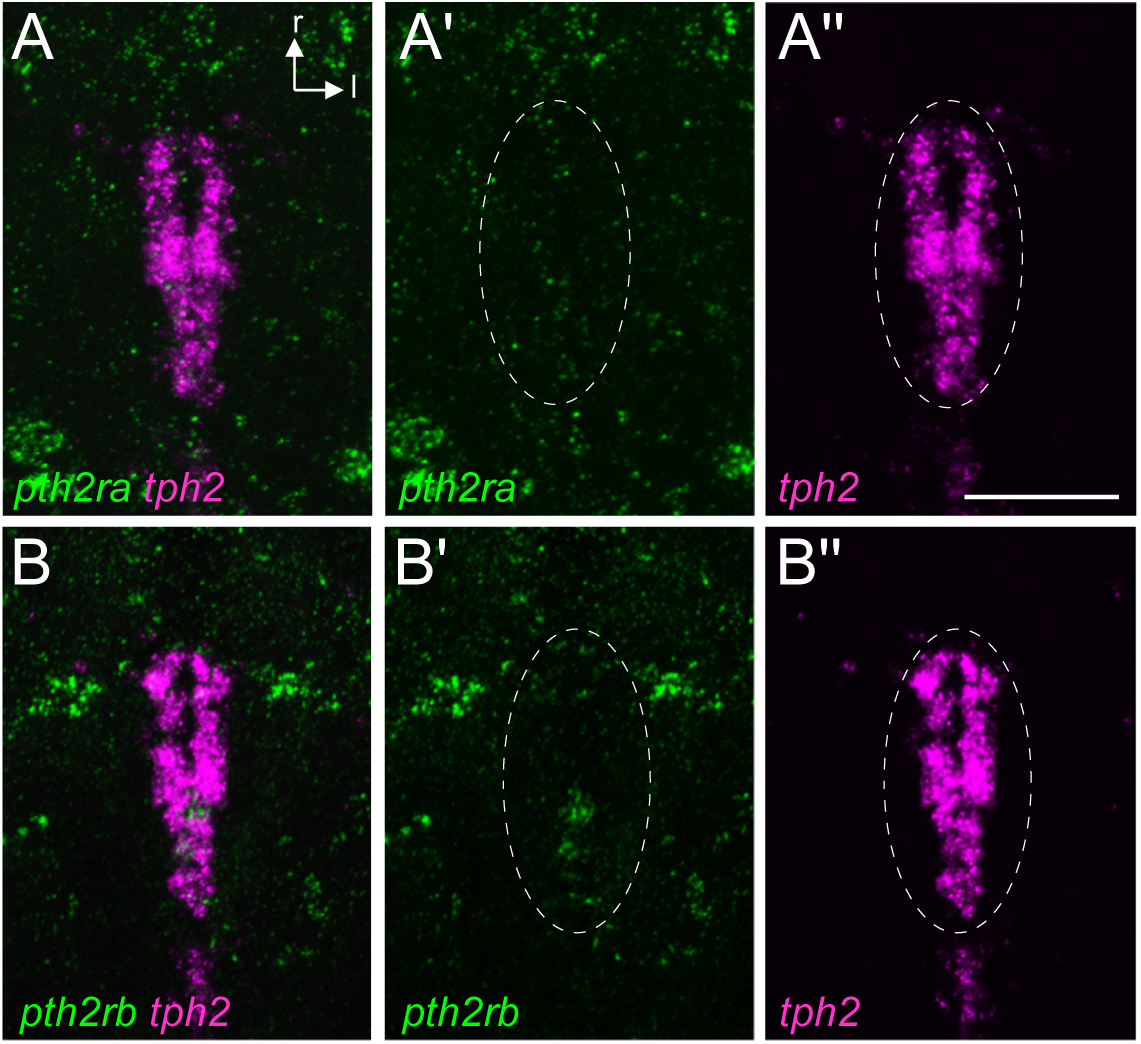
*pth2ra and pth2rb* are not expressed in raphe neurons. HCR using probes specific for *pth2ra* (**A,A’**), *pth2rb* (**B,B’**), and *tph2* (**A,A”,B,B”**) shows that these *pth receptors* are not co-expressed with *tph2*, which labels 5HT raphe neurons. Dashed circles indicate region of *tph2*-expressing 5HT raphe neurons. Scale bar: 50 μm. r, rostral; l, lateral.

**Figure S5.**
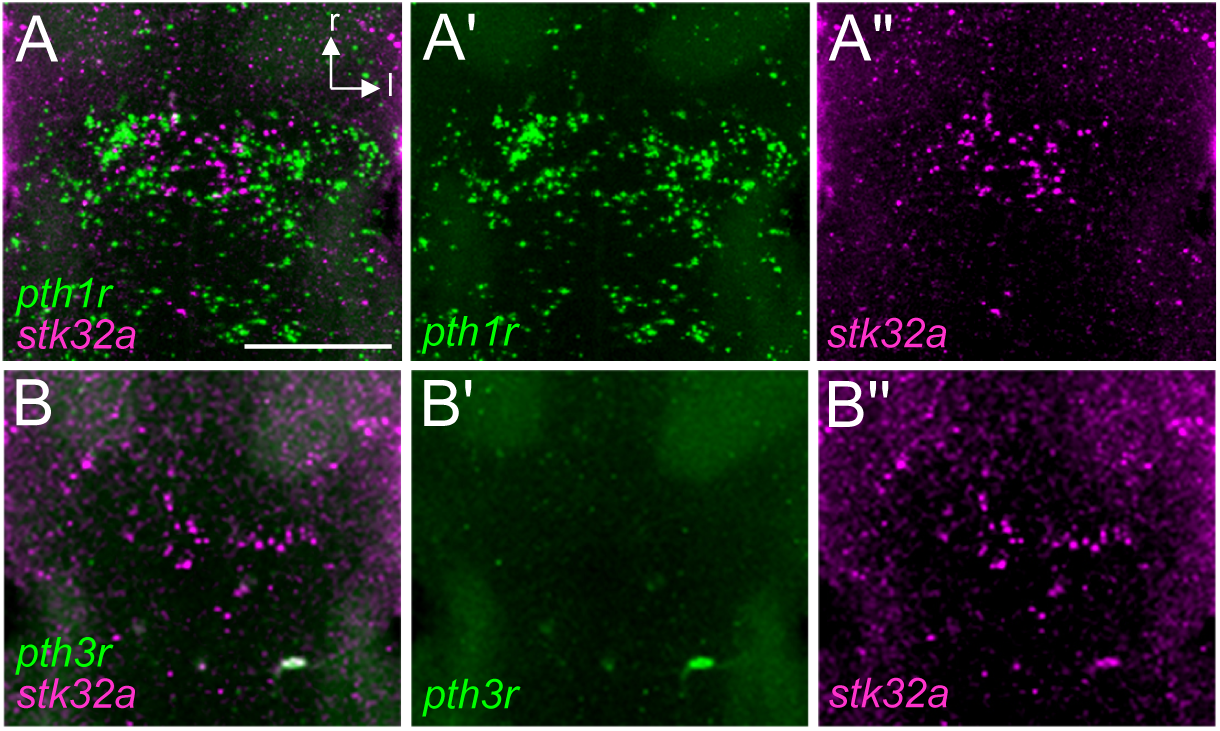
*pth1r* and *pth3r* are not expressed in prethalamic *stk32a* neurons. HCR using probes specific for *pth1r* (**A,A’**), *pth3r* (**B,B’**), and *stk32a* (**A,A”,B,B”**) shows that these *pth receptors* are not co-expressed with *stk32a* in the prethalamus. Scale bar: 50 μm. r, rostral; l, lateral.

**Video S1. Rotation showing QRFP-labeled and Hcrt-labeled neurons.** The video shows a rotation of the stack of confocal images shown in Figures 1B and 1B’ of a 5 dpf *Tg(qrfp:EGFP); Tg(hcrt:mRFP)* larval zebrafish brain.

**Video S2. Rotation showing QRFP-labeled and NPVF-labeled neurons.** The video shows a rotation of the stack of confocal images shown in Figures 1D and 1D’ of a 5 dpf *Tg(qrfp:EGFP); Tg(npvf:mCherry)* larval zebrafish brain.

## REFERENCES

1. Sehgal, A., and Mignot, E. (2011). Genetics of sleep and sleep disorders. Cell 146, 194–207. 10.1016/j.cell.2011.07.004.

2. Saper, C.B., and Fuller, P.M. (2017). Wake–sleep circuitry: an overview. Current Opinion in Neurobiology 44, 186–192. 10.1016/j.conb.2017.03.021.

3. Scammell, T.E., Arrigoni, E., and Lipton, J.O. (2017). Neural Circuitry of Wakefulness and Sleep. Neuron 93, 747–765. 10.1016/j.neuron.2017.01.014.

4. Liu, D., and Dan, Y. (2019). A Motor Theory of Sleep-Wake Control: Arousal-Action Circuit. Annual Review of Neuroscience 42, 27–46. 10.1146/annurev-neuro-080317-061813.

5. von Economo, C. (1930). Sleep as a problem of localization. The Journal of Nervous and Mental Disease 71, 249–259.

6. Adamantidis, A.R., and de Lecea, L. (2023). Sleep and the hypothalamus. Science 382, 405– 412. 10.1126/science.adh8285.

7. Nishino, S., Ripley, B., Overeem, S., Lammers, G.J., and Mignot, E. (2000). Hypocretin (orexin) deficiency in human narcolepsy. Lancet 355, 39–40.

8. Peyron, C., Faraco, J., Rogers, W., Ripley, B., Overeem, S., Charnay, Y., Nevsimalova, S., Aldrich, M., Reynolds, D., Albin, R., et al. (2000). A mutation in a case of early onset narcolepsy and a generalized absence of hypocretin peptides in human narcoleptic brains. Nat Med 6, 991–997.

9. de Lecea, L., Kilduff, T.S., Peyron, C., Gao, X., Foye, P.E., Danielson, P.E., Fukuhara, C., Battenberg, E.L., Gautvik, V.T., Bartlett, F.S., et al. (1998). The hypocretins: hypothalamus-specific peptides with neuroexcitatory activity. Proc Natl Acad Sci U S A 95, 322–327.

10. Chemelli, R.M., Willie, J.T., Sinton, C.M., Elmquist, J.K., Scammell, T., Lee, C., Richardson, J.A., Williams, S.C., Xiong, Y., Kisanuki, Y., et al. (1999). Narcolepsy in orexin knockout mice: molecular genetics of sleep regulation. Cell 98, 437–451.

11. Kaslin, J., Nystedt, J.M., Ostergard, M., Peitsaro, N., and Panula, P. (2004). The orexin/hypocretin system in zebrafish is connected to the aminergic and cholinergic systems. J Neurosci 24, 2678–2689.

12. Faraco, J.H., Appelbaum, L., Marin, W., Gaus, S.E., Mourrain, P., and Mignot, E. (2006). Regulation of hypocretin (orexin) expression in embryonic zebrafish. J Biol Chem 281, 29753–29761.

13. Prober, D.A., Rihel, J., Onah, A.A., Sung, R.J., and Schier, A.F. (2006). Hypocretin/orexin overexpression induces an insomnia-like phenotype in zebrafish. J Neurosci 26, 13400– 13410. 10.1523/JNEUROSCI.4332-06.2006.

14. Peyron, C., Sapin, E., Leger, L., Luppi, P.H., and Fort, P. (2009). Role of the melanin-concentrating hormone neuropeptide in sleep regulation. Peptides 30, 2052–2059. 10.1016/j.peptides.2009.07.022.

15. Jego, S., Glasgow, S.D., Herrera, C.G., Ekstrand, M., Reed, S.J., Boyce, R., Friedman, J., Burdakov, D., and Adamantidis, A.R. (2013). Optogenetic identification of a rapid eye movement sleep modulatory circuit in the hypothalamus. Nat Neurosci 16, 1637–1643. 10.1038/nn.3522.

16. Konadhode, R.R., Pelluru, D., Blanco-Centurion, C., Zayachkivsky, A., Liu, M., Uhde, T., Glen, W.B., van den Pol, A.N., Mulholland, P.J., and Shiromani, P.J. (2013). Optogenetic stimulation of MCH neurons increases sleep. J Neurosci 33, 10257–10263. 10.1523/JNEUROSCI.1225-13.2013.

17. Tsunematsu, T., Ueno, T., Tabuchi, S., Inutsuka, A., Tanaka, K.F., Hasuwa, H., Kilduff, T.S., Terao, A., and Yamanaka, A. (2014). Optogenetic manipulation of activity and temporally controlled cell-specific ablation reveal a role for MCH neurons in sleep/wake regulation. J Neurosci 34, 6896–6909. 10.1523/JNEUROSCI.5344-13.2014.

18. Vetrivelan, R., Kong, D., Ferrari, L.L., Arrigoni, E., Madara, J.C., Bandaru, S.S., Lowell, B.B., Lu, J., and Saper, C.B. (2016). Melanin-concentrating hormone neurons specifically promote rapid eye movement sleep in mice. Neuroscience 336, 102–113. 10.1016/j.neuroscience.2016.08.046.

19. Leung, L.C., Wang, G.X., Madelaine, R., Skariah, G., Kawakami, K., Deisseroth, K., Urban, A.E., and Mourrain, P. (2019). Neural signatures of sleep in zebrafish. Nature 571, 198– 204. 10.1038/s41586-019-1336-7.

20. Lee, D.A., Andreev, A., Truong, T.V., Chen, A., Hill, A.J., Oikonomou, G., Pham, U., Hong, Y.K., Tran, S., Glass, L., et al. (2017). Genetic and neuronal regulation of sleep by neuropeptide VF. eLife 6. 10.7554/eLife.25727.

21. Lee, D.A., Oikonomou, G., Cammidge, T., Andreev, A., Hong, Y., Hurley, H., and Prober, D.A. (2020). Neuropeptide VF neurons promote sleep via the serotonergic raphe. eLife 9, e54491. 10.7554/eLife.54491.

22. Barlow, I.L., and Rihel, J. (2017). Zebrafish sleep: from geneZZZ to neuronZZZ. Curr. Opin. Neurobiol. 44, 65–71. 10.1016/j.conb.2017.02.009.

23. Levitas-Djerbi, T., and Appelbaum, L. (2017). Modeling sleep and neuropsychiatric disorders in zebrafish. Curr. Opin. Neurobiol. 44, 89–93. 10.1016/j.conb.2017.02.017.

24. Oikonomou, G., and Prober, D.A. (2017). Attacking sleep from a new angle: contributions from zebrafish. Curr Opin Neurobiol 44, 80–88. 10.1016/j.conb.2017.03.009.

25. Stewart, A.M., Ullmann, J.F.P., Norton, W.H.J., Parker, M.O., Brennan, C.H., Gerlai, R., and Kalueff, A.V. (2015). Molecular psychiatry of zebrafish. Mol Psychiatry 20, 2–17. 10.1038/mp.2014.128.

26. Burgess, H.A., and Burton, E.A. (2023). A Critical Review of Zebrafish Neurological Disease Models−1. The Premise: Neuroanatomical, Cellular and Genetic Homology and Experimental Tractability. Oxf Open Neurosci 2, kvac018. 10.1093/oons/kvac018.

27. Rihel, J., Prober, D.A., Arvanites, A., Lam, K., Zimmerman, S., Jang, S., Haggarty, S.J., Kokel, D., Rubin, L.L., Peterson, R.T., et al. (2010). Zebrafish behavioral profiling links drugs to biological targets and rest/wake regulation. Science 327, 348–351. https://www.science.org/doi/10.1126/science.1183090.

28. Chiu, C.N., Rihel, J., Lee, D.A., Singh, C., Mosser, E.A., Chen, S., Sapin, V., Pham, U., Engle, J., Niles, B.J., et al. (2016). A Zebrafish Genetic Screen Identifies Neuromedin U as a Regulator of Sleep/Wake States. Neuron 89, 842–856. 10.1016/j.neuron.2016.01.007.

29. Renier, C., Faraco, J.H., Bourgin, P., Motley, T., Bonaventure, P., Rosa, F., and Mignot, E. (2007). Genomic and functional conservation of sedative-hypnotic targets in the zebrafish. Pharmacogenet Genomics 17, 237–253. 10.1097/FPC.0b013e3280119d62.

30. Elbaz, I., Yelin-Bekerman, L., Nicenboim, J., Vatine, G., and Appelbaum, L. (2012). Genetic ablation of hypocretin neurons alters behavioral state transitions in zebrafish. J Neurosci 32, 12961–12972. 10.1523/JNEUROSCI.1284-12.2012.

31. Singh, C., Oikonomou, G., and Prober, D.A. (2015). Norepinephrine is required to promote wakefulness and for hypocretin-induced arousal in zebrafish. eLife 4, e07000. 10.7554/eLife.07000.

32. Oikonomou, G., Altermatt, M., Zhang, R., Coughlin, G.M., Montz, C., Gradinaru, V., and Prober, D.A. (2019). The Serotonergic Raphe Promote Sleep in Zebrafish and Mice. Neuron 103, 686–701.e8. 10.1016/j.neuron.2019.05.038.

33. Liu, J., Merkle, F.T., Gandhi, A.V., Gagnon, J.A., Woods, I.G., Chiu, C.N., Shimogori, T., Schier, A.F., and Prober, D.A. (2015). Evolutionarily conserved regulation of hypocretin neuron specification by Lhx9. Development 142, 1113–1124. 10.1242/dev.117424.

34. Chen, A., Chiu, C.N., Mosser, E.A., Kahn, S., Spence, R., and Prober, D.A. (2016). QRFP and Its Receptors Regulate Locomotor Activity and Sleep in Zebrafish. J Neurosci 36, 1823–1840. 10.1523/JNEUROSCI.2579-15.2016.

35. Chartrel, N., Alonzeau, J., Alexandre, D., Jeandel, L., Alvear-Perez, R., Leprince, J., Boutin, J., Vaudry, H., Anouar, Y., and Llorens-Cortes, C. (2011). The RFamide neuropeptide 26RFa and its role in the control of neuroendocrine functions. Frontiers in neuroendocrinology 32, 387–397. 10.1016/j.yfrne.2011.04.001.

36. Mirabeau, O., and Joly, J.-S. (2013). Molecular evolution of peptidergic signaling systems in bilaterians. Proceedings of the National Academy of Sciences 110, E2028–E2037. 10.1073/pnas.1219956110.

37. Primeaux, S.D., Barnes, M.J., and Braymer, H.D. (2013). Hypothalamic QRFP: Regulation of Food Intake and Fat Selection. Horm Metab Res 45, 967–974. 10.1055/s-0033-1353181.

38. Ukena, K., Osugi, T., Leprince, J., Vaudry, H., and Tsutsui, K. (2014). Molecular evolution of GPCRs: 26Rfa/GPR103. Journal of molecular endocrinology 52, T119–31. 10.1530/JME-13-0207.

39. Xu, B., Bergqvist, C.A., Sundström, G., Lundell, I., Vaudry, H., Leprince, J., and Larhammar, D. (2015). Characterization of peptide QRFP (26RFa) and its receptor from amphioxus, *Branchiostoma floridae*. General and Comparative Endocrinology 210, 107– 113. 10.1016/j.ygcen.2014.10.010.

40. Leprince, J., Bagnol, D., Bureau, R., Fukusumi, S., Granata, R., Hinuma, S., Larhammar, D., Primeaux, S., Sopkova-de Oliveiras Santos, J., Tsutsui, K., et al. (2017). The Arg–Phe-amide peptide 26RFa/glutamine RF-amide peptide and its receptor: IUPHAR Review 24. British Journal of Pharmacology 174, 3573–3607. 10.1111/bph.13907.

41. Suarez-Bregua, P., Torres-Nuñez, E., Saxena, A., Guerreiro, P., Braasch, I., Prober, D.A., Moran, P., Cerda-Reverter, J.M., Du, S.J., Adrio, F., et al. (2016). Pth4, an ancient parathyroid hormone lost in eutherian mammals, reveals a new brain-to-bone signaling pathway. The FASEB Journal 31, 569–583. 10.1096/fj.201600815R.

42. Suarez-Bregua, P., Saxena, A., Bronner, M.E., and Rotllant, J. (2017). Targeted Pth4-expressing cell ablation impairs skeletal mineralization in zebrafish. PLOS ONE 12, e0186444. 10.1371/journal.pone.0186444.

43. Singh, C., Rihel, J., and Prober, D.A. (2017). Neuropeptide Y regulates sleep by modulating noradrenergic signaling. Curr Biol 27, 3796–3811.e5. 10.1016/j.cub.2017.11.018.

44. Weber, F., and Dan, Y. (2016). Circuit-based interrogation of sleep control. Nature 538, 51– 59. 10.1038/nature19773.

45. Madelaine, R., Lovett-Barron, M., Halluin, C., Andalman, A.S., Liang, J., Skariah, G.M., Leung, L.C., Burns, V.M., and Mourrain, P. (2017). The hypothalamic NPVF circuit modulates ventral raphe activity during nociception. Sci Rep 7, 41528. 10.1038/srep41528.

46. Sakurai, T., Amemiya, A., Ishii, M., Matsuzaki, I., Chemelli, R.M., Tanaka, H., Williams, S.C., Richardson, J.A., Kozlowski, G.P., Wilson, S., et al. (1998). Orexins and orexin receptors: a family of hypothalamic neuropeptides and G protein-coupled receptors that regulate feeding behavior. Cell 92, 573–585.

47. Takayasu, S., Sakurai, T., Iwasaki, S., Teranishi, H., Yamanaka, A., Williams, S.C., Iguchi, H., Kawasawa, Y.I., Ikeda, Y., Sakakibara, I., et al. (2006). A neuropeptide ligand of the G protein-coupled receptor GPR103 regulates feeding, behavioral arousal, and blood pressure in mice. Proc Natl Acad Sci U S A 103, 7438–7443.

48. Liu, Q., Guan, X.M., Martin, W.J., McDonald, T.P., Clements, M.K., Jiang, Q., Zeng, Z., Jacobson, M., Williams, D.L., Yu, H., et al. (2001). Identification and characterization of novel mammalian neuropeptide FF-like peptides that attenuate morphine-induced antinociception. J Biol Chem 276, 36961–36969. 10.1074/jbc.M105308200.

49. Lin, J.Y., Knutsen, P.M., Muller, A., Kleinfeld, D., and Tsien, R.Y. (2013). ReaChR: a red-shifted variant of channelrhodopsin enables deep transcranial optogenetic excitation. Nat Neurosci 16, 1499–1508. 10.1038/nn.3502.

50. Sharrock, A.V., Mulligan, T.S., Hall, K.R., Williams, E.M., White, D.T., Zhang, L., Emmerich, K., Matthews, F., Nimmagadda, S., Washington, S., et al. (2022). NTR 2.0: a rationally engineered prodrug-converting enzyme with substantially enhanced efficacy for targeted cell ablation. Nat Methods 19, 205–215. 10.1038/s41592-021-01364-4.

51. Carter, M.E., Brill, J., Bonnavion, P., Huguenard, J.R., Huerta, R., and de Lecea, L. (2012). Mechanism for Hypocretin-mediated sleep-to-wake transitions. Proc Natl Acad Sci U S A 109, E2635–44. 10.1073/pnas.1202526109.

52. Suarez-Bregua, P., Cal, L., Cañestro, C., and Rotllant, J. (2017). PTH Reloaded: A New Evolutionary Perspective. Frontiers in Physiology 8.

53. Rubin, D.A., and Jüppner, H. (1999). Zebrafish Express the Common Parathyroid Hormone/Parathyroid Hormone-related Peptide Receptor (PTH1R) and a Novel Receptor (PTH3R) That Is Preferentially Activated by Mammalian and Fugufish Parathyroid Hormone-related Peptide. Journal of Biological Chemistry 274, 28185–28190. 10.1074/jbc.274.40.28185.

54. Rubin, D.A., Hellman, P., Zon, L.I., Lobb, C.J., Bergwitz, C., and Jüppner, H. (1999). A G Protein-coupled Receptor from Zebrafish Is Activated by Human Parathyroid Hormone and Not by Human or Teleost Parathyroid Hormone-related Peptide: IMPLICATIONS FOR THE EVOLUTIONARY CONSERVATION OF CALCIUM-REGULATING PEPTIDE HORMONES. Journal of Biological Chemistry 274, 23035–23042. 10.1074/jbc.274.33.23035.

55. Meyer, M.P., and Smith, S.J. (2006). Evidence from in vivo imaging that synaptogenesis guides the growth and branching of axonal arbors by two distinct mechanisms. J Neurosci 26, 3604–3614. 10.1038/nn.2654.

56. Mathias, J.R., Zhang, Z., Saxena, M.T., and Mumm, J.S. (2014). Enhanced cell-specific ablation in zebrafish using a triple mutant of Escherichia coli nitroreductase. Zebrafish 11, 85–97. 10.1089/zeb.2013.0937.

57. Tabor, K.M., Bergeron, S.A., Horstick, E.J., Jordan, D.C., Aho, V., Porkka-Heiskanen, T., Haspel, G., and Burgess, H.A. (2014). Direct activation of the Mauthner cell by electric field pulses drives ultra-rapid escape responses. J Neurophysiol 112, 834–844. 10.1152/jn.00228.2014.

58. Hull, K.L., Fathimani, K., Sharma, P., and Harvey, S. (1998). Calcitropic peptides: neural perspectives. Comparative Biochemistry and Physiology Part C: Pharmacology, Toxicology and Endocrinology 119, 389–410. 10.1016/S0742-8413(98)00010-3.

59. Lanske, B., Karaplis, A.C., Lee, K., Luz, A., Vortkamp, A., Pirro, A., Karperien, M., Defize, L.H.K., Ho, C., Mulligan, R.C., et al. (1996). PTH/PTHrP Receptor in Early Development and Indian Hedgehog—Regulated Bone Growth. Science 273, 663–666. 10.1126/science.273.5275.663.

60. Miao, D., He, B., Karaplis, A.C., and Goltzman, D. (2002). Parathyroid hormone is essential for normal fetal bone formation. J Clin Invest 109, 1173–1182. 10.1172/JCI14817.

61. Kronenberg, H.M. (2006). PTHrP and Skeletal Development. Annals of the New York Academy of Sciences 1068, 1–13. 10.1196/annals.1346.002.

62. Usdin, T.B., Paciga, M., Riordan, T., Kuo, J., Parmelee, A., Petukova, G., Camerini-Otero, R.D., and Mezey, É. (2008). Tuberoinfundibular Peptide of 39 Residues Is Required for Germ Cell Development. Endocrinology 149, 4292–4300. 10.1210/en.2008-0419.

63. Kwong, R.W.M., and Perry, S.F. (2015). An Essential Role for Parathyroid Hormone in Gill Formation and Differentiation of Ion-Transporting Cells in Developing Zebrafish. Endocrinology 156, 2384–2394. 10.1210/en.2014-1968.

64. Anneser, L., Alcantara, I.C., Gemmer, A., Mirkes, K., Ryu, S., and Schuman, E.M. (2020). The neuropeptide Pth2 dynamically senses others via mechanosensation. Nature 588, 653–657. 10.1038/s41586-020-2988-z.

65. Anneser, L., Gemmer, A., Eilers, T., Alcantara, I.C., Loos, A.-Y., Ryu, S., and Schuman, E.M. (2022). The neuropeptide Pth2 modulates social behavior and anxiety in zebrafish. iScience 25, 103868. 10.1016/j.isci.2022.103868.

66. Keller, D., Láng, T., Cservenák, M., Puska, G., Barna, J., Csillag, V., Farkas, I., Zelena, D., Dóra, F., Küppers, S., et al. (2022). A thalamo-preoptic pathway promotes social grooming in rodents. Current Biology 32, 4593–4606.e8. 10.1016/j.cub.2022.08.062.

67. Sun, J., Yuan, Y., Wu, X., Liu, A., Wang, J., Yang, S., Liu, B., Kong, Y., Wang, L., Zhang, K., et al. (2022). Excitatory SST neurons in the medial paralemniscal nucleus control repetitive self-grooming and encode reward. Neuron 110, 3356–3373.e8. 10.1016/j.neuron.2022.08.010.

68. Kripke, D.F., Lavie, P., Parker, D., Huey, L., and Deftos, L.J. (1978). Plasma Parathyroid Hormone and Calcium Are Related to Sleep Stage Cycles*. The Journal of Clinical Endocrinology & Metabolism 47, 1021–1027. 10.1210/jcem-47-5-1021.

69. Shafer, M.E.R., Sawh, A.N., and Schier, A.F. (2022). Gene family evolution underlies cell-type diversification in the hypothalamus of teleosts. Nat Ecol Evol 6, 63–76. 10.1038/s41559-021-01580-3.

70. Steuernagel, L., Lam, B.Y.H., Klemm, P., Dowsett, G.K.C., Bauder, C.A., Tadross, J.A., Hitschfeld, T.S., del Rio Martin, A., Chen, W., de Solis, A.J., et al. (2022). HypoMap—a unified single-cell gene expression atlas of the murine hypothalamus. Nat Metab 4, 1402– 1419. 10.1038/s42255-022-00657-y.

71. Herb, B.R., Glover, H.J., Bhaduri, A., Colantuoni, C., Bale, T.L., Siletti, K., Hodge, R., Lein, E., Kriegstein, A.R., Doege, C.A., et al. (2023). Single-cell genomics reveals region-specific developmental trajectories underlying neuronal diversity in the human hypothalamus. Science Advances 9, eadf6251. 10.1126/sciadv.adf6251.

72. Arrigoni, E., Chee, M.J.S., and Fuller, P.M. (2019). To eat or to sleep: That is a lateral hypothalamic question. Neuropharmacology 154, 34–49. 10.1016/j.neuropharm.2018.11.017.

73. Clift, D., Richendrfer, H., Thorn, R.J., Colwill, R.M., and Creton, R. (2014). High-Throughput Analysis of Behavior in Zebrafish Larvae: Effects of Feeding. Zebrafish 11, 455–461. 10.1089/zeb.2014.0989.

74. Aharon, D., and Marlow, F.L. (2021). Sexual determination in zebrafish. Cell. Mol. Life Sci. 79, 8. 10.1007/s00018-021-04066-4.

75. Dreosti, E., Lopes, G., Kampff, A.R., and Wilson, S.W. (2015). Development of social behavior in young zebrafish. Front Neural Circuits 9, 39. 10.3389/fncir.2015.00039.

76. Chen, S., Chiu, C.N., McArthur, K.L., Fetcho, J.R., and Prober, D.A. (2016). TRP channel mediated neuronal activation and ablation in freely behaving zebrafish. Nature Methods 13, 147–150. 10.1038/nmeth.3691.

77. Chen, A., Singh, C., Oikonomou, G., and Prober, D.A. (2017). Genetic Analysis of Histamine Signaling in Larval Zebrafish Sleep. eNeuro 4. 10.1523/ENEURO.0286-16.2017.

78. Hunsley, M.S., and Palmiter, R.D. (2003). Norepinephrine-deficient mice exhibit normal sleep-wake states but have shorter sleep latency after mild stress and low doses of amphetamine. Sleep 26, 521–526.

79. Thakkar, M.M. (2011). Histamine in the regulation of wakefulness. Sleep Med Rev 15, 65– 74. 10.1016/j.smrv.2010.06.004.

80. Berridge, C.W., Schmeichel, B.E., and Espana, R.A. (2012). Noradrenergic modulation of wakefulness/arousal. Sleep Med Rev 16, 187–197. 10.1016/j.smrv.2011.12.003.

81. Venner, A., Broadhurst, R.Y., Sohn, L.T., Todd, W.D., and Fuller, P.M. (2020). Selective activation of serotoninergic dorsal raphe neurons facilitates sleep through anxiolysis. Sleep 43. 10.1093/sleep/zsz231.

82. Chen, S., Reichert, S., Singh, C., Oikonomou, G., Rihel, J., and Prober, D.A. (2017). Light-dependent regulation of sleep/wake states by prokineticin 2 in zebrafish. Neuron 95, 153–168.e6. 10.1016/j.neuron.2017.06.001.

83. Robles, E., Laurell, E., and Baier, H. (2014). The Retinal Projectome Reveals Brain-Area-Specific Visual Representations Generated by Ganglion Cell Diversity. Current Biology 24, 2085–2096. 10.1016/j.cub.2014.07.080.

84. Takahashi, T.M., Sunagawa, G.A., Soya, S., Abe, M., Sakurai, K., Ishikawa, K., Yanagisawa, M., Hama, H., Hasegawa, E., Miyawaki, A., et al. (2020). A discrete neuronal circuit induces a hibernation-like state in rodents. Nature 583, 109–114. 10.1038/s41586-020-2163-6.

85. Romanov, R.A., Zeisel, A., Bakker, J., Girach, F., Hellysaz, A., Tomer, R., Alpár, A., Mulder, J., Clotman, F., Keimpema, E., et al. (2017). Molecular interrogation of hypothalamic organization reveals distinct dopamine neuronal subtypes. Nat Neurosci 20, 176–188. 10.1038/nn.4462.

86. Seifinejad, A., Ramosaj, M., Shan, L., Li, S., Possovre, M.-L., Pfister, C., Fronczek, R., Garrett-Sinha, L.A., Frieser, D., Honda, M., et al. (2023). Epigenetic silencing of selected hypothalamic neuropeptides in narcolepsy with cataplexy. Proceedings of the National Academy of Sciences 120, e2220911120. 10.1073/pnas.2220911120.

87. Okamoto, K., Yamasaki, M., Takao, K., Soya, S., Iwasaki, M., Sasaki, K., Magoori, K., Sakakibara, I., Miyakawa, T., Mieda, M., et al. (2016). QRFP-Deficient Mice Are Hypophagic, Lean, Hypoactive and Exhibit Increased Anxiety-Like Behavior. PLOS ONE 11, e0164716. 10.1371/journal.pone.0164716.

88. Li, W., Yuan, L., Tong, G., He, Y., Meng, Y., Hao, S., Chen, J., Guo, J., Bringhurst, R., and Yang, D. (2018). Phospholipase C signaling activated by parathyroid hormone mediates the rapid osteoclastogenesis in the fracture healing of orchiectomized mice. BMC Musculoskeletal Disorders 19, 311. 10.1186/s12891-018-2231-3.

89. Moen, M.D., and Scott, L.J. (2006). Recombinant Full-Length Parathyroid Hormone (1–84). Drugs 66, 2371–2381. 10.2165/00003495-200666180-00008.

90. Cosman, F., Nieves, J.W., and Dempster, D.W. (2017). Treatment Sequence Matters: Anabolic and Antiresorptive Therapy for Osteoporosis. Journal of Bone and Mineral Research 32, 198–202. 10.1002/jbmr.3051.

91. Everson, C.A., Folley, A.E., and Toth, J.M. (2012). Chronically Inadequate Sleep Results in Abnormal Bone Formation and Abnormal Bone Marrow in Rats. Exp Biol Med (Maywood) 237, 1101–1109. 10.1258/ebm.2012.012043.

92. Xu, X., Wang, L., Chen, L., Su, T., Zhang, Y., Wang, T., Ma, W., Yang, F., Zhai, W., Xie, Y., et al. (2016). Effects of chronic sleep deprivation on bone mass and bone metabolism in rats. Journal of Orthopaedic Surgery and Research 11, 87. 10.1186/s13018-016-0418-6.

93. Swanson, C.M., Shea, S.A., Wolfe, P., Cain, S.W., Munch, M., Vujović, N., Czeisler, C.A., Buxton, O.M., and Orwoll, E.S. (2017). Bone Turnover Markers After Sleep Restriction and Circadian Disruption: A Mechanism for Sleep-Related Bone Loss in Humans. The Journal of Clinical Endocrinology & Metabolism 102, 3722–3730. 10.1210/jc.2017-01147.

94. Swanson, C.M., Kohrt, W.M., Buxton, O.M., Everson, C.A., Wright, K.P., Orwoll, E.S., and Shea, S.A. (2018). The importance of the circadian system & sleep for bone health. Metabolism 84, 28–43. 10.1016/j.metabol.2017.12.002.

95. Staab, J.S., Smith, T.J., Wilson, M., Montain, S.J., and Gaffney-Stomberg, E. (2019). Bone turnover is altered during 72 h of sleep restriction: a controlled laboratory study. Endocrine 65, 192–199. 10.1007/s12020-019-01937-6.

96. Fu, X., Zhao, X., Lu, H., Jiang, F., Ma, X., and Zhu, S. (2011). Association between sleep duration and bone mineral density in Chinese women. Bone 49, 1062–1066. 10.1016/j.bone.2011.08.008.

97. Kobayashi, D., Takahashi, O., Deshpande, G.A., Shimbo, T., and Fukui, T. (2012). Association between osteoporosis and sleep duration in healthy middle-aged and elderly adults: a large-scale, cross-sectional study in Japan. Sleep Breath 16, 579–583. 10.1007/s11325-011-0545-6.

98. Tian, Y., Shen, L., Wu, J., Xu, G., Yang, S., Song, L., Zhang, Y., Mandiwa, C., Yang, H., Liang, Y., et al. (2015). Sleep duration and timing in relation to osteoporosis in an elderly Chinese population: a cross-sectional analysis in the Dongfeng–Tongji cohort study. Osteoporos Int 26, 2641–2648. 10.1007/s00198-015-3172-4.

99. Wang, K., Wu, Y., Yang, Y., Chen, J., Zhang, D., Hu, Y., Liu, Z., Xu, J., Shen, Q., Zhang, N., et al. (2015). The associations of bedtime, nocturnal, and daytime sleep duration with bone mineral density in pre- and post-menopausal women. Endocrine 49, 538–548. 10.1007/s12020-014-0493-6.

100. Sun, Y., Tisdale, R.K., and Kilduff, T.S. (2021). Hypocretin/Orexin Receptor Pharmacology and Sleep Phases. The Orexin System. Basic Science and Role in Sleep Pathology 45, 22–37. 10.1159/000514963.

101. Agetsuma, M., Aizawa, H., Aoki, T., Nakayama, R., Takahoko, M., Goto, M., Sassa, T., Amo, R., Shiraki, T., Kawakami, K., et al. (2010). The habenula is crucial for experience-dependent modification of fear responses in zebrafish. Nat Neurosci 13, 1354–1356. 10.1038/nn.2654.

102. Chen, S., Oikonomou, G., Chiu, C.N., Niles, B.J., Liu, J., Lee, D.A., Antoshechkin, I., and Prober, D.A. (2013). A large-scale in vivo analysis reveals that TALENs are significantly more mutagenic than ZFNs generated using context-dependent assembly. Nucleic Acids Res 41, 2769–2778. 10.1093/nar/gks1356.

103. Schneider, C.A., Rasband, W.S., and Eliceiri, K.W. (2012). NIH Image to ImageJ: 25 years of image analysis. Nature Methods 9, 671–675.

104. Labun, K., Montague, T.G., Gagnon, J.A., Thyme, S.B., and Valen, E. (2016). CHOPCHOP v2: a web tool for the next generation of CRISPR genome engineering. Nucleic Acids Res. 10.1093/nar/gkw398.

105. Griesbeck, O., Baird, G.S., Campbell, R.E., Zacharias, D.A., and Tsien, R.Y. (2001). Reducing the Environmental Sensitivity of Yellow Fluorescent Protein. Mechanism and applications. Journal of Biological Chemistry 276, 29188–29194. 10.1074/jbc.M102815200.

106. Distel, M., Wullimann, M.F., and Koster, R.W. (2009). Optimized Gal4 genetics for permanent gene expression mapping in zebrafish. Proc Natl Acad Sci U S A 106, 13365– 13370. 10.1073/pnas.0903060106.

107. Bergeron, S.A., Hannan, M.C., Codore, H., Fero, K., Li, G.H., Moak, Z., Yokogawa, T., and Burgess, H.A. (2012). Brain selective transgene expression in zebrafish using an NRSE derived motif. Front Neural Circuits 6, 110. 10.3389/fncir.2012.00110.

108. Xie, X., Mathias, J.R., Smith, M.A., Walker, S.L., Teng, Y., Distel, M., Koster, R.W., Sirotkin, H.I., Saxena, M.T., and Mumm, J.S. (2013). Silencer-delimited transgenesis: NRSE/RE1 sequences promote neural-specific transgene expression in a NRSF/REST-dependent manner. BMC Biol 10, 93. 10.1186/1741-7007-10-93.

109. Kawakami, K. (2007). Tol2: a versatile gene transfer vector in vertebrates. Genome Biology 8, S7. 10.1186/gb-2007-8-s1-s7.

110. Lee, D.A., Oikonomou, G., and Prober, D.A. (2022). Large-scale Analysis of Sleep in Zebrafish. Bio-protocol 12, e4313. 10.21769/BioProtoc.4313.

111. Lauter, G., Söll, I., and Hauptmann, G. (2011). Multicolor fluorescent in situ hybridization to define abutting and overlapping gene expression in the embryonic zebrafish brain. Neural Dev 6, 10. 10.1186/1749-8104-6-10.

112. Choi, H.M.T., Schwarzkopf, M., Fornace, M.E., Acharya, A., Artavanis, G., Stegmaier, J., Cunha, A., and Pierce, N.A. (2018). Third-generation in situ hybridization chain reaction: multiplexed, quantitative, sensitive, versatile, robust. Development 145, dev165753. 10.1242/dev.165753.

